# The MNK1/2-eIF4E axis drives melanoma plasticity, progression, and resistance to immunotherapy

**DOI:** 10.1101/2020.05.29.117531

**Authors:** Fan Huang, Christophe Gonçalves, Margarita Bartish, Joelle Rémy-Sarrazin, Qianyu Guo, Audrey Emond, William Yang, Dany Plourde, Jie Su, Marina Godoy Gimeno, Yao Zhan, Mikhael Attias, Alba Galán, Tomasz Rzymski, Milena Mazan, Magdalena Masiejczyk, Jacek Faber, Elie Khoury, Alexandre Benoit, Natascha Gagnon, David Dankort, Ciriaco A. Piccirillo, Fabrice Journe, Ghanem Ghanem, H. Uri Saragovi, Nahum Sonenberg, Ivan Topisirovic, Wilson H. Miller, Sonia V. del Rincón

## Abstract

Melanomas commonly undergo a phenotype switch, from a proliferative to an invasive state. Melanoma plasticity exhibited as phenotype switching contributes to immunotherapy resistance, however the mechanisms are not completely understood and thus therapeutically unexploited. Here, using a transgenic melanoma mouse model, we demonstrated a critical role of the MNK1/2-eIF4E axis in melanoma plasticity and resistance to immunotherapy. We showed that phospho-eIF4E deficient murine melanomas express high levels of melanocytic antigens, with similar results verified in patient melanomas. Mechanistically, we identified that phospho-eIF4E controls the translation of *NGFR*, a critical effector of phenotype switching. In patients with melanoma, the expression of *MKNK1*, the kinase for eIF4E, positively correlated with markers of immune exhaustion. Genetic ablation of phospho-eIF4E reprogrammed the immunosuppressive microenvironment, exemplified by lowered production of inflammatory factors and increased CD8^+^ T cell infiltrates. Blocking phospho-eIF4E, using MNK1/2 inhibitors, offers a new strategy to inhibit melanoma plasticity and improve the survival response to anti-PD-1 immunotherapy.

## Introduction

Malignant melanoma is the deadliest form of skin cancer. In melanoma, the major signaling pathways, RAS/RAF/MAPK and PI3K/AKT, are constitutively activated through numerous avenues, including genetic alterations in BRAF and PTEN, respectively. Both pathways ultimately converge upon the eukaryotic translation initiation factor 4E (eIF4E) to induce its phosphorylation. For instance, *BRAF^V600E^* is the most common activating mutation in cutaneous melanoma and is upstream of mitogen-activated protein kinase (MAPK)-interacting serine/threonine-protein kinase 1 (MNK1) and 2 (MNK2), which directly phosphorylate eIF4E^1–3^. Loss of function mutations in *PTEN*, occurring in up to 30% of patients with *BRAF* mutant melanoma, on the other hand, will indirectly result in the phosphorylation of eIF4E via activation of the mTOR-4E-BP1 axis^1^. Many groups, including our own, have shown that dysregulated mRNA translation by aberrant activation of the MNK1/2-eIF4E axis, plays a critical role in tumor progression to invasive and metastatic disease^4–6^. Moreover, the phosphorylation of eIF4E is increased during melanoma progression^7,8^. The phosphorylation of eIF4E on serine 209 (S209), catalyzed exclusively by MNK1 and MNK2^2,3^, increases the oncogenic potential of eIF4E^7–9^. Mechanistically, phospho-eIF4E selectively increases the translation of a subset of mRNAs encoding proteins involved in cell survival, proliferation and metastasis^10–12^.

Immunotherapies blocking immune checkpoints, such as CTLA-4 and the PD-1/PD-L1 axis, show therapeutic efficacy in patients with metastatic melanoma^13–16^. Antibodies against CTLA-4, PD-1 or PD-L1 are currently approved for clinical use, or are in clinical trials, in many types of cancers^17–19^. Despite the approval of immunotherapy for the management of a growing number of cancer types, checkpoint blockade as monotherapy has achieved limited clinical success in most malignancies. Many patients fail to respond, and others, who initially had responded, eventually experience tumor relapse^19,20^. Therapeutic efficacy of combination immune checkpoint inhibitors against CTLA-4 and PD-1 can also be hampered by the appearance of sometimes life-threatening immune-related adverse events^18,21,22^. Key factors that favour a response to immune checkpoint blockade include the presence of immune cell infiltration and the availability of tumor-associated target antigens^23–27^. Melanoma plasticity, exhibited as phenotype switching, has been proposed to contribute to primary or secondary resistance to immunotherapy and is a process akin to the epithelial-mesenchymal-like transition (EMT)^28–35^. During phenotype switching, melanomas undergo dedifferentiation with (1) a loss of melanocytic antigens, (2) increased pro-inflammatory cytokine/chemokine secretion, and (3) tumor infiltration of myeloid-derived suppressor cells (MDSCs)^28–31,33–38^. However, the mechanisms underlying the regulation of the phenotype switch are not completely understood, and thus, few therapeutic approaches have been exploited to prevent or block this process^28^.

Herein, we investigated the role of phospho-eIF4E in melanoma progression and tumor immunity. We hypothesized that phospho-eIF4E promotes melanoma phenotype switching, metastasis, and functions also in non-melanoma cells to support an immune suppressive microenvironment that is unfavorable for an effective response to immunotherapy. MNK1/2 inhibitors, blocking the phosphorylation of eIF4E, therefore serve a dual purpose as (1) therapies that inhibit tumor plasticity and (2) immune-modulatory agents that can be used in combination with immunotherapies in metastatic melanoma.

## Results

### Phospho-eIF4E promotes melanoma outgrowth and metastasis

To determine whether phospho-eIF4E contributes to melanoma development and metastasis *in vivo*, we crossed eIF4E^S209A/S209A^ mice, wherein eIF4E cannot be phosphorylated^9^, with the well-described *Tyr::CreER*/*BRaf^CA/+^/Pten^lox/lox^* conditional melanoma model^39,40^. This melanoma model allows 4-hydroxytamoxifen (4-HT) inducible melanocyte-targeted *BRAF^V600E^* expression and simultaneous *PTEN* inactivation (Figure 1A). As expected, following topical administration of 4-HT on the lower back of mice, hyperpigmented lesions were observed within 12-15 days and melanomas developed within 2-3 weeks (Figure 1B and S1A). Compared to the *BRaf^CA/+^/Pten^lox/lox^* (referred to hereafter as eIF4E^WT^) mice, the melanoma outgrowth in *Braf^CA/+^/Pten^lox/lox^/Eif4e^S209A/S209A^* (henceforth termed eIF4E^KI^) mice, devoid of phosphorylated eIF4E, was robustly decreased (Figure 1B and 1C). While we did not observe any significant difference in tumor initiation between the eIF4E^WT^ and eIF4E^KI^ mice (Figure S1A), Ki67 staining revealed a less proliferative state of the eIF4E^KI^ melanomas (Figure 1D), indicating that phospho-eIF4E promotes melanoma proliferation *in vivo*. The protection against primary melanoma outgrowth is dependent on the dosage of phospho-eIF4E loss, as mice expressing one copy of eIF4E and one copy of eIF4E S209A (eIF4E^HET^) exhibited a delay in melanoma outgrowth compared to the eIF4E^WT^ mice (Figure S1B). Importantly, eIF4E^KI^ mice exhibit a significant increase in survival compared to their eIF4E^WT^ counterparts (Figure 1E). Using ultrasound, we detected an acute enlargement of the regional/draining lymph nodes (i.e. inguinal lymph nodes) in the eIF4E^KI^ mice, but not the eIF4E^WT^ mice, at day 12 after 4-HT administration (Figure 1F). H&E staining revealed no metastasis at this time point (data not shown), suggesting an acute inflammatory response rather than early metastasis in these lymph nodes^41,42^. On day 24, while there was no significant difference in inguinal lymph node size between the two cohorts of mice, we detected regional lymph node metastasis in the eIF4E^WT^ mice but not the eIF4E^KI^ mice (Figure 1F and S1C). As the primary melanoma progressed, we observed significantly accelerated enlargement of the draining lymph nodes in the eIF4E^WT^ mice compared to the eIF4E^KI^ mice (Figure 1F). Metastasis to the distant cervical lymph nodes was assessed when primary tumors were size matched between 500 mm^3^ to 800 mm^3^. Numbers of metastasis-positive cervical lymph nodes were significantly lower in the eIF4E^KI^ mice compared to the eIF4E^WT^ mice (Figure 1G). These results suggest that inhibiting the phosphorylation of eIF4E profoundly hinders melanoma outgrowth and metastasis.

**Figure 1.**
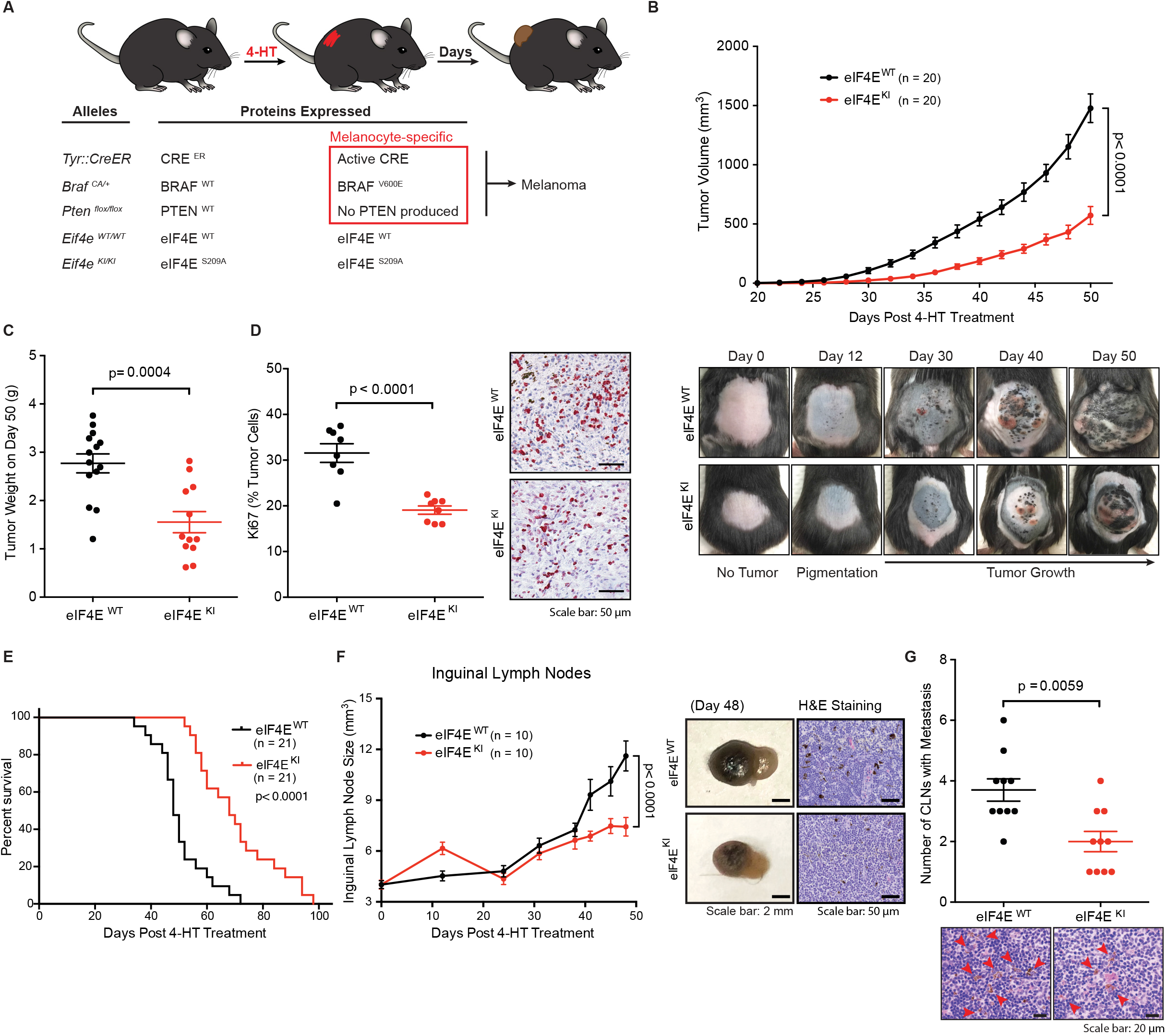
Phospho-eIF4E-deficiency decreases melanoma outgrowth and metastasis. **(A)** Schematic diagram of the *Tyr::CreER/BRaf*^*CA*/+^/*Pten*^*flox/flox*^/*Eif4e*^*WT/WT*^ (eIF4E^WT^) and the *Tyr::CreER/BRaf*^*CA*^/+/*Pten*^*flox/flox*^/*Eif4e^KI/KI^* (eIF4E^KI^) murine melanoma model. **(B)** Tumor growth curve (left) and representative pictures (right) of eIF4E^WT^ and eIF4E^KI^ mice (n = 20 per group) after topical administration of 4-HT. Two-way ANOVA. **(C)** Primary tumor weight (Day 50) of eIF4E^WT^ (n = 14) and eIF4E^KI^ (n = 12) mice. Two-sided unpaired t test. **(D)** Percentages (left) and representative images (right) of Ki67-positve melanoma cells in eIF4E^WT^ and eIF4E^KI^ primary melanoma sections (Day 50, n = 12 per genotype). Two-sided unpaired t test. **(E)** Kaplan–Meier curves showing overall survival of eIF4E^WT^ and eIF4E^KI^ mice (n = 21 per group). Log-rank test. **(F)** Inguinal lymph node (ILN) sizes measured by ultrasound (left), representative ILN pictures (Day 48, middle) and representative images of H&E-stained ILNs (Day 48, right) of eIF4E^WT^ and eIF4E^KI^ mice (n = 10 mice per group). Two-way ANOVA. **(G)** Cervical lymph nodes (CLNs) were resected from eIF4E^WT^ and eIF4E^KI^ mice (n = 10 mice per group) with size-matched primary tumors (500-800 mm3). Number of metastasis-positive CLNs per mouse (left) and representative images of H&E-stained CLNs (right) are presented. Mann Whitney test. All values are represented as mean ± SEM.

### Inhibition of phospho-eIF4E blocks melanoma dedifferentiation and loss of melanocytic antigens

We next characterized the histology of melanomas derived from eIF4E^WT^ and eIF4E^KI^ transgenic mice. Upon inspection of the H&E stained primary melanomas, while no significant morphological differences were observed, we noted a striking increase in melanin pigmentation in the eIF4E^KI^ melanomas, as compared to eIF4E^WT^ melanomas (Figure 2A and S2A). In addition, immunohistochemical staining for the melanoma marker S100 showed that the frequency of pigmented S100-positive melanoma cells is significantly higher in the eIF4E^KI^ cohort, compared to the eIF4E^WT^ cohort (Figure 2B). Melanin pigmentation is tightly controlled by the microphthalmia-associated transcription factor (MITF), which is critical for the survival of pigmented cells and drives melanocyte differentiation^43^. Dedifferentiated melanomas, characterized by a loss, or low expression, of MITF, have been associated with increased invasion and metastasis, resistance to chemotherapies and immunotherapies, and reduced overall patient survival^29,30^. eIF4E^KI^ melanomas express more MITF, compared to eIF4E^WT^ melanomas (Figure 2C). Moreover, the expression of two melanocytic differentiation antigens, Melan-A (also known as MART-1) and GP100 (also known as Pmel17)^44,45^, are significantly increased in the eIF4E^KI^ melanomas (Figure 2D). Consistent with previous studies^40,46,47^, Melan-A expression is virtually undetectable in most of the eIF4E^WT^ melanomas (Figure 2D and S2B). Melanomas harboring a 50% reduction in the phosphorylation of eIF4E (eIF4E^HET^) also showed increased Melan-A expression, relative to eIF4E^WT^ melanomas (Figure S2B). In support of the clinical relevance of our findings, the expression of phospho-eIF4E is negatively correlated with the expression of Melan-A in patient-derived melanomas (Figure 2E and S2C). Melanomas with high overall expression of phospho-eIF4E have significantly lower expression, or, in many patient samples, no expression of Melan-A. Conversely, melanomas with low overall phospho-eIF4E expression show intense Melan-A staining (Figure 2E). Intriguingly, in patient samples with regional phospho-eIF4E expression, the phospho-eIF4E staining was mutually exclusive with Melan-A expression (Figure S2C). Together, these data suggest that repressing the phosphorylation of eIF4E in melanoma results in a differentiated phenotype characterized by the retention of pigmentation and melanocytic antigens.

**Figure 2.**
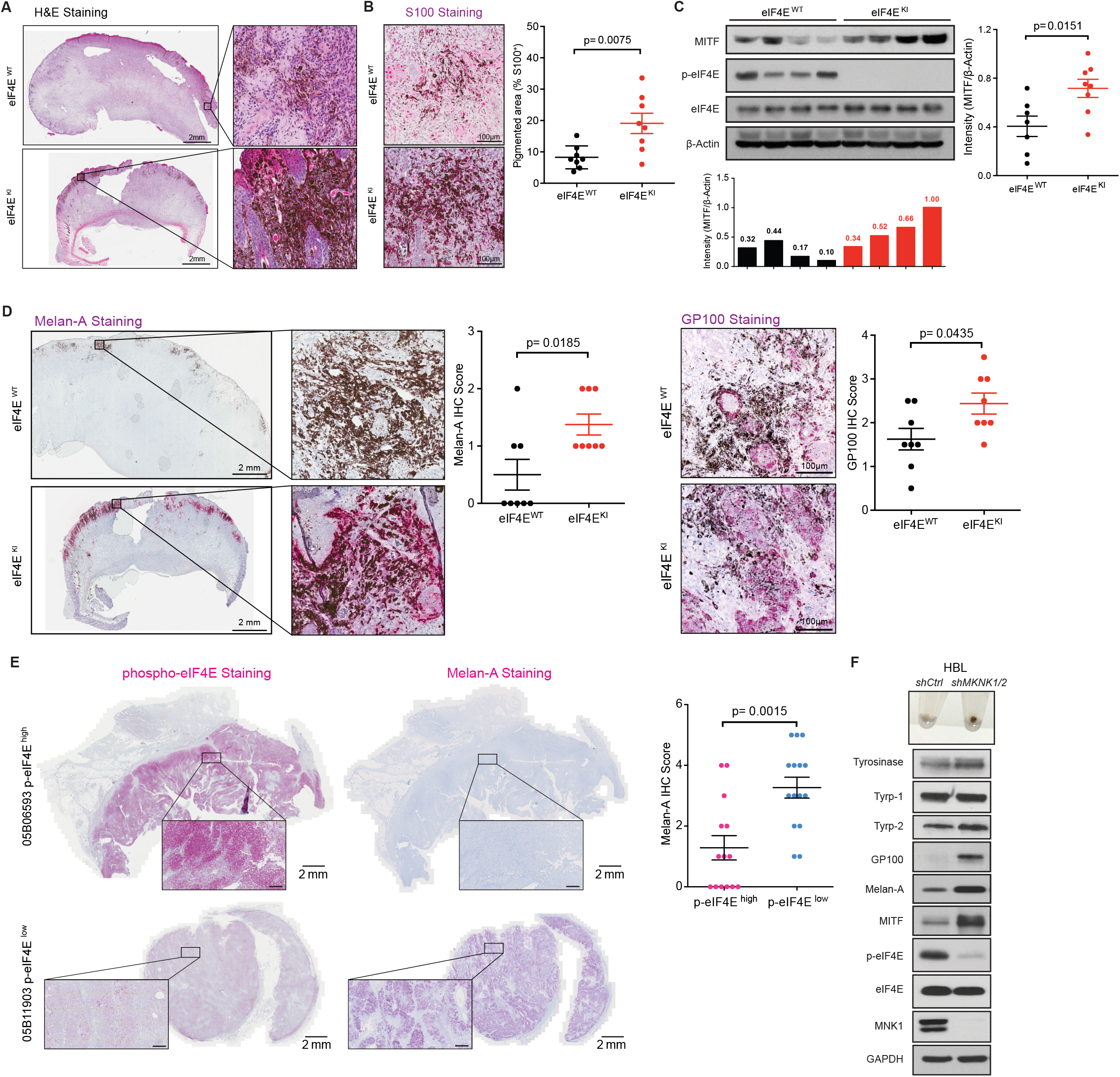
Phospho-eIF4E deficient murine and human melanomas are more differentiated. **(A)** Representative eIF4E^WT^ (n = 16) and eIF4E^KI^ (n = 15) primary tumor histology with H&E staining (Day 50). **(B)** Representative IHC images showing the expression of S100 in eIF4E^WT^ and eIF4E^KI^ melanomas (left), and the percentage of pigmented area in the S100-positive region in the eIF4E^WT^ and eIF4E^KI^ tumors (Day 50, n = 8 per group). Two-sided unpaired t test. **(C)** Representative western blot of the indicated proteins in 4 eIF4E^WT^ (from a total n = 7) and 4 eIF4E^KI^ (from a total n = 8) primary melanomas (left top, Day 50) with quantification of MITF normalized to β-Actin level (left bottom and right). Two-sided unpaired t test. **(D)** Representative IHC images with scores showing the expression of Melan-A (left) and GP100 (right) in eIF4E^WT^ and eIF4E^KI^ melanomas (Day 50, n = 8 per group). Mann Whitney test. **(E)** Left panel: IHC images showing the expression of phospho-eIF4E and Melan-A in 2 representative tumors from a total of 29 patients with melanoma (top, patient with high overall phospho-eIF4E expression and bottom, patient with low overall phospho-eIF4E expression). Right panel: IHC scoring of Melan-A in phospho-eIF4E^high^ (n = 14) and phospho-eIF4E^low^ (n = 15) patient-derived melanomas. Mann Whitney test. **(F)** Western blot analysis of the indicated proteins in human HBL melanoma cells stably expressing *shCtrl* and *shMKNK1/2*. Results are representative of n = 3 independent experiments. All values are represented as mean ± SEM.

We next reasoned that blocking the activity of MNK1/2, the kinases that phosphorylate eIF4E, would reverse tumor cell plasticity in human melanoma cells. To test this, we used the invasive patient-derived melanoma cell line HBL wherein we previously knocked down *MKNK1* and *MKNK2* using shRNA^48^. *MKNK1/2* knockdown in HBL cells showed reduced phospho-eIF4E level concomitant with increased pigmentation (Figure 2F). Silencing of *MKNK1/2* in HBL cells also resulted in an increased expression of MITF and induction of the melanogenic proteins tyrosinase, Tyrp-2, Melan-A and GP100, as compared to their shRNA control counterparts (Figure 2F). Together, these data suggest that inhibition of the MNK1/2-eIF4E axis reverses melanoma plasticity toward a differentiated phenotype.

### Phospho-eIF4E deficient melanomas are less invasive through an inhibition of NGFR

Melanomas that have undergone dedifferentiation, or phenotype switching, are characterized by a more invasive phenotype^29,38,49–55^. As we have shown that the mice harboring more differentiated eIF4E^KI^ melanomas also have decreased lymph node metastasis *in vivo*, we next wanted to recapitulate this phenotype *in vitro*. We derived murine melanoma cell lines, MDMel-WT (No. 73 and 88) and MDMel-KI (No. 58 and 61), from two eIF4E^WT^ and two eIF4E^KI^ tumor-bearing animals, respectively. All cell lines were confirmed as PTEN-negative (*Pten^lox/lox^*), *Tyr::CreER*-positive and *Braf^CA/+^*, similar to D4M.3a murine melanoma cells previously derived from the eIF4E^WT^ mouse model^46^ (Figure 3A, S3A and S3F). Characterization of these cell lines revealed distinct phenotypes. The MDMel-KI tumor derived cell lines, which are phospho-eIF4E deficient, failed to invade as efficiently as the MDMel-WT cells (Figure 3B and Figure S3B), despite being more proliferative *in vitro* (Figure S3C and S3D). Notably, previous studies by our group showed that the *MKNK1/2* knockdown HBL cells are less invasive compared to their *shCtrl* counterparts, similar to the phospho-eIF4E-deficient MDMel-KI phenotype^48^.

**Figure 3.**
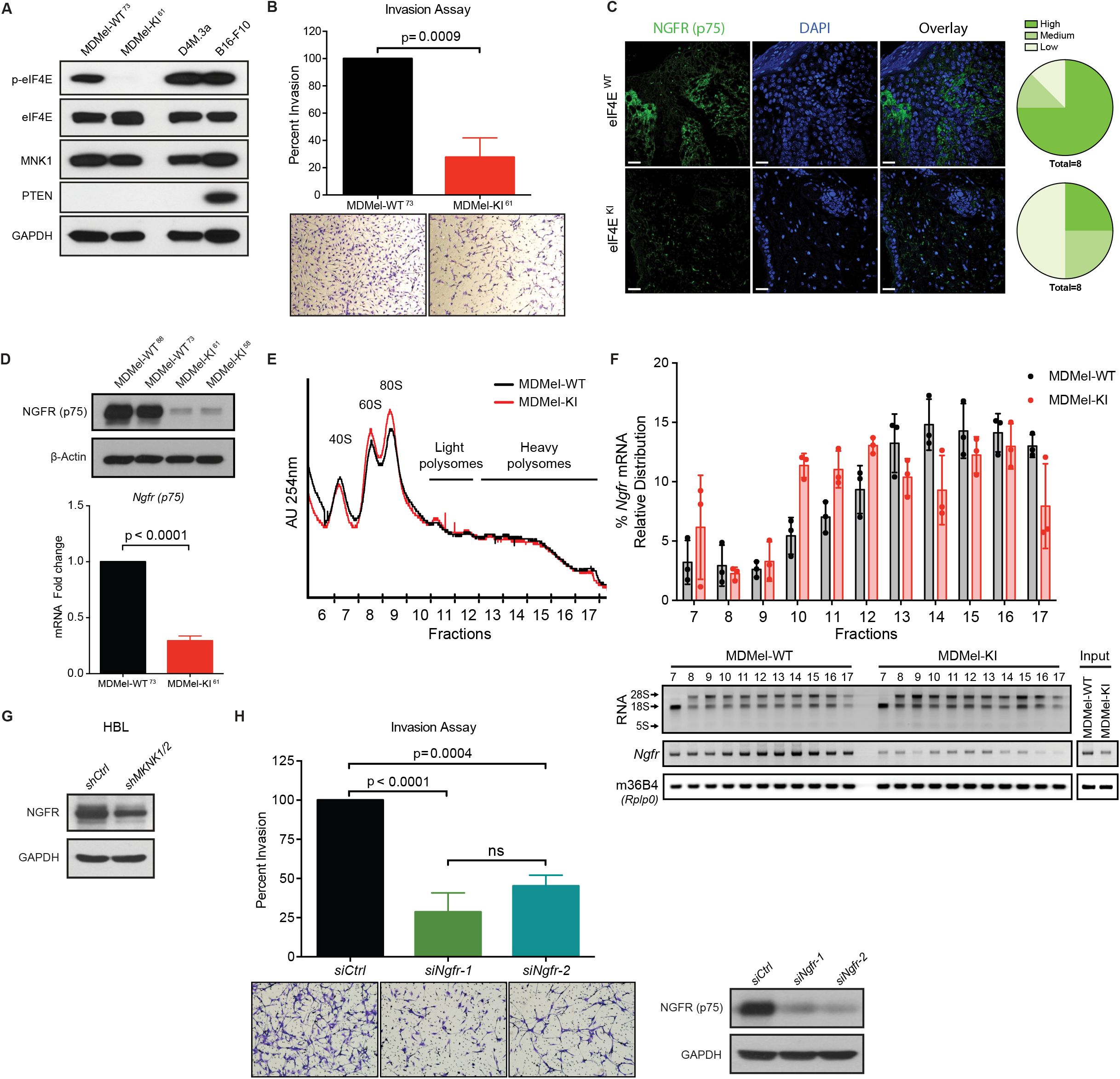
Phospho-eIF4E deficient melanoma cells resist phenotype switching via inhibiting NGFR mRNA translation. **(A)** Western blot analysis of the indicated proteins in murine melanoma cell lines MDMel-WT^73^, MDMel-KI^61^, D4M.3a and B16-F10 (representative of n = 3). **(B)** Percent invasion of MDMel-KI^61^ cells relative to MDMel-WT^73^ cells (top) and representative images with original magnification (×10, bottom). Results are representative of n = 3 independent experiments. Two-sided unpaired t-test. **(C)** Representative immunofluorescence images showing the expression of NGFR in eIF4E^WT^ and eIF4E^KI^ tumors (Day 50, n = 8 per group) (left) and the percentage of samples with high-, medium- and low-expression of NGFR from each group (right). **(D)** Top: Western blot analysis of NGFR expression in the murine melanoma cell lines MDMel-WT^88^, MDMel-WT^73^, MDMel-KI^61^ and MDMel-KI^58^ (representative of n = 3). Bottom: Percentage of *Ngfr* mRNA in MDMel-KI^61^ cells relative to MDMel-WT^73^ cells, normalized to m36B4 (*Rplp0*) as a reference gene (n = 4). Two-sided unpaired t-test. See also Figure S3F. **(E)** Polysome profiles of MDMel-WT and MDMel-KI cells (representative of n = 3). **(F)** Percentage of transcripts in each polysomal fraction quantified by RT-qPCR (top) and representative image showing ribosomal RNAs and PCR-amplified cDNA fragments of the indicated targets (bottom). Values represent the mean ± SD (n = 3). **(G)** Western blot of NGFR expression in human HBL melanoma cells stably expressing *shCtrl* and *shMKNK1/2* (representative of n = 3). **(H)** Left panel: Percent invasion of MDMel-WT^73^ cells with *Ngfr* knockdown relative to the control group (top), and representative images with original magnification (×10, bottom, see also Figure S3G). Right panel: Western blot confirming knockdown of NGFR. Results are representative of n = 3 independent experiments. Two-sided unpaired t-test. All values are represented as mean ± SD.

The data presented thus far support a previously unappreciated role for phospho-eIF4E in regulating melanoma plasticty exhibited as dedifferentiation/phenotype switching. A wealth of data suggests that the cell surface receptor NGFR (p75) serves as a molecular switch to promote melanoma dedifferentiation^31,33,36,37^. We show that eIF4E^KI^ melanomas have significantly less NGFR expression compared to the eIF4E^WT^ tumors (Figure 3C). Consistent with previous findings^31,37^, the regions staining positive for NGFR are restricted to areas that are negative for the melanosomal marker Melan-A (Figure S3E). In addition, the expression of NGFR was significantly decreased in both MDMel-KI melanoma cell lines, compared to the two MDMel-WT cell lines (Figure 3D and S3F). Although the *Ngfr* mRNA was decreased by about 3-5 fold in the MDMel-KI cells (Figure 3D and S3F), given the robust decrease of NGFR at protein level and the crucial role of phospho-eIF4E in mRNA translation, we next determined whether NGFR expression could be under the translational control of phospho-eIF4E. We subjected our MDMel cell lines to polysome profiling, a technique used to separate efficiently translated mRNAs bound to heavy polysomes from poorly translated mRNAs in light polysomes^56,57^. Representative polysome gradient profiles from MDMel-KI and MDMel-WT samples overlap (Figure 3E), consistent with the role of phospho-eIF4E in regulating the translation of selective mRNAs, without exerting major effects on global protein synthesis^9,10,48,58^. Quantitative PCR analysis of RNA isolated from heavy and light polysome-bound fractions in MDMel-WT and MDMel-KI cells indicated that a lack of eIF4E phosphorylation leads to a redistribution of *Ngfr* mRNAs from heavy (efficiently translated) to light (poorly translated) polysomes (Figure 3F). This is consistent with the notion that the phosphorylation of eIF4E bolsters the translation of *Ngfr* mRNA. Control over the regulation of NGFR expression, upstream of phospho-eIF4E, is MNK1/2 dependent, as the expression of NGFR protein is inhibited in *MKNK1/2* silenced HBL cells (Figure 3G). Furthermore, knockdown of *Ngfr* in the MDMel-WT cells significantly impaired cell invasion (Figure 3H), with no change in proliferation (Figure S3G and S3H). Together, our data suggest that repressing the MNK1/2-eIF4E axis suppresses melanoma dedifferentiation and invasion by decreasing NGFR protein expression, a key effector of phenotype switching^31,33,36,37^.

### Inhibition of phospho-eIF4E blocks pro-inflammatory cytokine secretion

Inflammation promotes dedifferentiation in melanoma, which in turn leads to an altered cytokine/chemokine profile^30,33,38,59,60^. Thus, we next monitored the expression of chemokines and cytokines present in the conditioned media harvested from eIF4E^WT^ and eIF4E^KI^ melanomas (Figure 4A). A set of secreted factors that are known to promote cancer cell invasion were found to be decreased in the eIF4E^KI^ melanomas. These included Angiopoietin-2 (ANGPT2), CCL2, CCL12, CCL5, IGFBP-2, IGFBP-6, IL-6 and MMP-9 (Figure 4B and S4A). Interestingly, many of these factors (i.e. Angiopoietins, CCL2, CCL12, IGFBP-2, IGFBP-6) have never been thoroughly investigated in melanoma phenotype switching, but are represented in large scale RNAseq analyses of genes associated with phenotype switching^53–55,61,62^. Moreover, among these cytokines, many have been associated with the expansion (such as IL-6), recruitment (such as CCL2, CCL12, and CCL5) and function (such as MMP-9) of immune suppressive cells, such as myeloid-derived suppressor cells (MDCSs)^63–66^. Given the importance of CCL5 and IL-6 in inflammation and immune suppression, we used ELISA to further validate that these two factors were significantly repressed in the supernatant harvested from the eIF4E^KI^ melanomas compared with the eIF4E^WT^ melanomas (Figure 4C).

**Figure 4.**
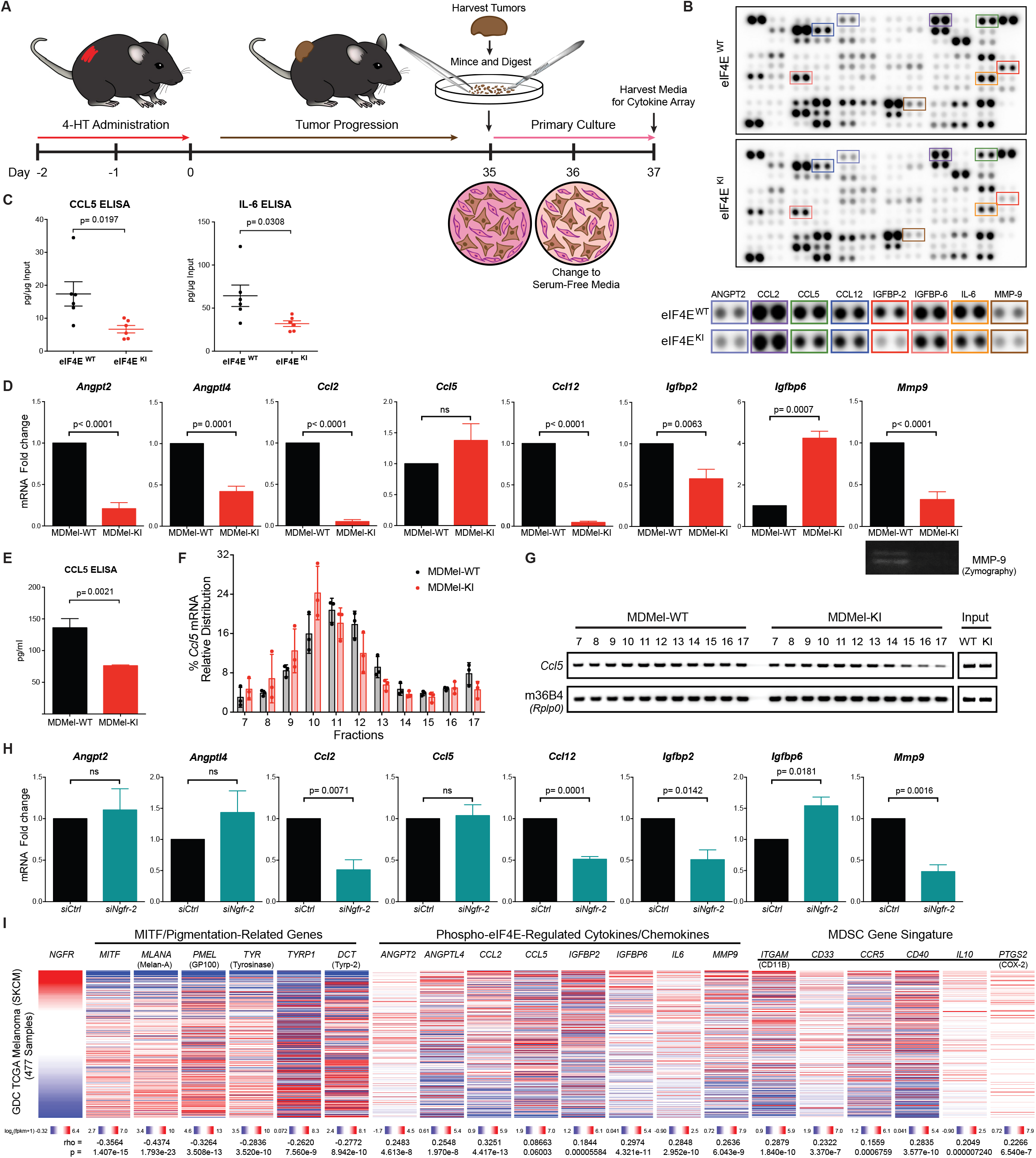
Phospho-eIF4E deficient melanomas have an altered secretome. **(A)** Schematic of the experimental design for the membrane-based cytokine arrays. **(B)** Representative images of the cytokine arrays showing the expression of secreted factors present in the conditioned medium derived from eIF4E^WT^ and eIF4E^KI^ primary melanoma cultures (n = 2 mice per genotype). **(C)** Concentration of CCL5 and IL-6 in the conditioned medium derived from the eIF4E^WT^ and eIF4E^KI^ primary melanoma cultures (n = 6 mice per genotype). Data are represented as mean ± SEM. Two-sided unpaired t-test. **(D)** Percentage of the indicated mRNAs in MDMel-KI cells relative to MDMel-WT cells, normalized to m36B4 (*Rplp0*) as a reference gene (n = 3 for *Igfbp6*, n = 5 for the rest). Two-sided unpaired t-test. Zymography to assess MMP-9 activity in the conditioned medium of MDMel-WT and MDMel-KI cells (bottom, representative of n = 3). **(E)** Concentration of CCL5 in the conditioned medium of MDMel-WT and MDMel-KI cells. Values represent the mean ± SD (n=3). Two-sided unpaired t-test. **(F)** Percentage of transcripts in each polysomal fraction quantified by RT-qPCR. Values represent the mean ± SD (n = 3). **(G)** Representative image showing PCR-amplified cDNA fragments of the indicated targets (n = 3). **(H)** Percentage of indicated mRNAs in *siNgfr-2*-transfected MDMel-WT cells relative to the control group, normalized to m36B4 (*Rplp0*) as a reference gene. Values represent the mean ± SD (n = 3). Two-sided unpaired t-test. **(I)** Correlation of the expression of indicated genes (HTSeq - FPKM) with *NGFR* expression (HTSeq - FPKM) in GDC TCGA Melanoma dataset (SKCM, n = 477). Spearman rank-order.

We next sought to uncover the molecular mechanisms underlying the differential expression of soluble factors from the eIF4E^WT^ and eIF4E^KI^ melanomas. First, we used our murine derived melanoma cell lines, devoid or not of phospho-eIF4E, to interrogate the cellular source of the differentially regulated factors. While the expression of most of these factors are decreased at the mRNA level (Figure 4D and S4B), CCL5 was decreased only at the protein level in the MDMel-KI cells, with no significant change in *Ccl5* mRNA compared to the MDMel-WT cells (Figure 4D and 4E). Similar to our results in MDMel cell lines, mouse embryonic fibroblasts (MEFs) devoid of phospho-eIF4E (KI MEFs)^9^ express less CCL5 protein by ELISA (Figure S4C), although *Ccl5* mRNA levels remain unchanged compared to WT MEFs (Figure S4E). Analysis of *Ccl5* mRNAs in the polysomal fractions isolated from the MDMel-KI cells showed that there was a significant shift of *Ccl5* mRNAs towards the translationally less active polysome fractions (Figure 4F and 4G). Similar polysome profiling results revealed a strong shift of *Ccl5* mRNAs from heavy-to light-polysome fractions in KI MEFs compared to WT MEFs (Figure S4D and S4E). Our data thus demonstrate that *Ccl5* mRNA is a novel translational target of phospho-eIF4E.

Although IL-6 mRNA and protein were not detectable *in vitro* in MDMel cell lines or MEFs (data not shown), we detected IL-6 in the primary tumors derived from the MDMel-WT or MDMel-KI cells injected into C57BL/6 mice (Figure S4F). Consistent with the cytokine array result, we detected a robust decrease in IL-6 mRNA and protein expression in the MDMel-KI-derived melanoma, compared to their MDMel-WT counterparts (Figure S4F). An up-regulation in IL-6 expression is consistent with melanoma phenotype switching^37,67^. Our results provide evidence that IL-6 expression is induced by the interaction of the tumor cells with the tumor microenvironment (TME) and is dependent on the phosphorylation status of eIF4E in the tumor cells.

As we have shown that phospho-eIF4E promotes melanoma phenotype switching via enhancing the translation of *Ngfr*, we next asked whether the expression of the phospho-eIF4E-regulated cytokine/chemokines were also dependent on the expression of NGFR. To test this, we knocked down *Ngfr* in MDMel-WT cells using siRNA (*siNgfr-2*) (Figure 3H and S4G). Strikingly, the changes of cytokine/chemokine mRNA profile following *Ngfr* knockdown resembled those of the MDMel-KI cells. Such changes included the down-regulation of *Ccl2*, *Ccl12*, *Igfbp2, Mmp9*, and an unexpected up-regulation of *Igfbp6* (Figure 4H). *Angpt2* and *Angptl4* remain unchanged, indicating that the expression of these two genes may not be directly regulated through NGFR (further discussed in Figure 7 and Discussion).

We examined TCGA data for the relationship between the differentiation/pigmentation gene set (*MITF*, *Mlana*, *PMEL*, *TYR*, *TYRP1* and *DCT*) and *NGFR* expression. As expected, the expression of *NGFR* in human melanomas correlated negatively with the differentiation/pigmentation gene set (Figure 4I, left panel)^31,36,37^. Furthermore, *NGFR* expression also positively correlated with the phospho-eIF4E-regulated cytokine/chemokine gene signature (*ANGPT2*, *ANGPTL4*, *CCL2*, *IGFBP2* and *MMP9*) (Figure 4I, middle panel), and genes frequently associated with myeloid derived suppressor cells (MDSC) influx (*ITGAM*, *CD33*, *CCR5*, *CD40*, *IL10*, and *PTGS2*) (Figure 4I, right panel)^63–66^. Notably, *NGFR* gene expression does not correlate with *CCL5* mRNA in the TCGA melanoma cohort (Figure 4I, middle panel), further supporting that CCL5 is under translational control by phospho-eIF4E.

### Phospho-eIF4E supports an immune suppressive tumor microenvironment

eIF4E^KI^ melanomas show evidence of resisting phenotype switching, with (1) decreased NGFR expression, (2) repressed production of pro-inflammatory secreted factors and (3) maintenance of Melan-A antigen expression, leading us to predict that suppressing phospho-eIF4E favors anti-tumor immunity. BRAF^V600E^/PTEN^null/null^ murine melanomas have been characterized by the presence immune suppressive MDSCs and modest T cell infiltration^35,68^. Thus, this model can be used to assess the impact of melanomas lacking phospho-eIF4E on tumor immune cell infiltration. Immune phenotyping of eIF4E^WT^ and eIF4E^KI^ melanomas showed that the phospho-eIF4E deficient melanomas were significantly more infiltrated with CD3^+^ T-cell and CD8^+^ cytotoxic T-cell populations, compared to eIF4E^WT^ melanomas (Figure 5A, S5A, S5C and S5D). Importantly, the eIF4E^KI^ melanomas showed higher intra-tumoral Granzyme B-positive cell infiltrates, a marker of cytolytic action (Figure 5B). Consistent with the pro-inflammatory cytokine signature associated with eIF4E^WT^ melanomas, these melanomas were more infiltrated with MDSCs, compared to eIF4E^KI^ melanomas (Figure 5A, S5B and S5C). There were significantly fewer monocytic-MDSCs in the eIF4E^KI^ melanomas, compared to their control counterparts (Figure 5A and S5C). Statistical analysis revealed that tumor outgrowth in our model is negatively correlated with the numbers of tumor-infiltrating T cells and positively correlated with the numbers of tumor-infiltrating MDSCs (Figure S5C). No significant changes in dendritic cells or macrophages were found between the eIF4E^WT^ and eIF4E^KI^ melanomas (Figure S5E). Overall, these data suggest an anti-tumor immune effect in the melanomas where eIF4E can not be phosphorylated.

**Figure 5.**
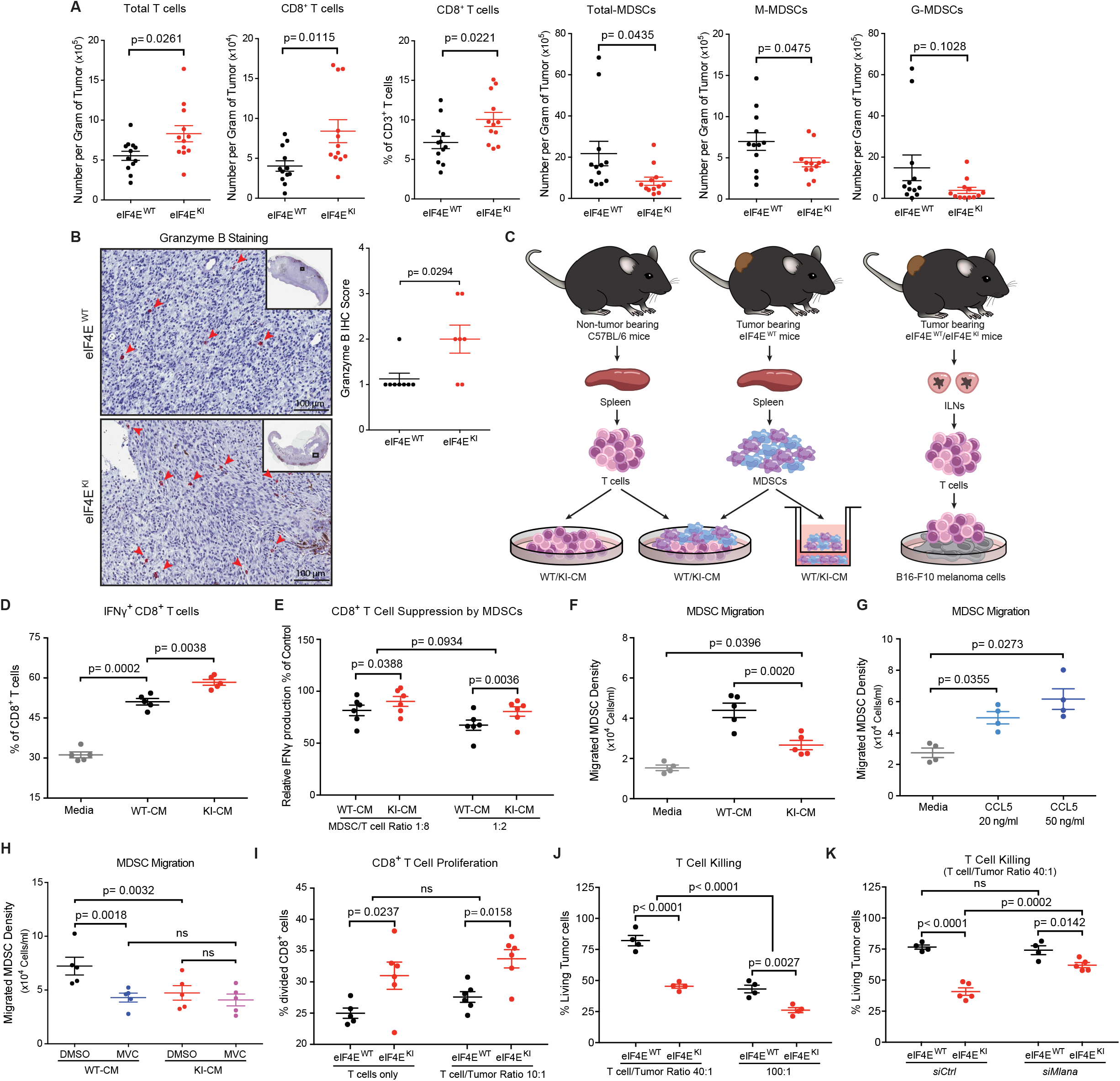
Phospho-eIF4E-deficiency impairs melanoma immunosuppression. **(A)** Immune cell populations infiltrated into the melanomas from eIF4E^WT^ and eIF4E^KI^ mice (Day 50, n = 12 per group). Two-sided unpaired t-test. **(B)** Representative eIF4E^WT^ (n = 8) and eIF4E^KI^ (n = 7) tumor samples (Day 50) with IHC staining of Granzyme B (left) and corresponding scores (right). Mann Whitney test. **(C)** Schematic of *ex vivo* experimental designs. Left: T cell activation, MDSC/T cell co-culture and MDSC migration assay. Right: T cell/Tumor cell co-culture assay. **(D)** Percentage of CD8^+^ T cells producing IFNγ, following stimulation with CD3/CD28 (72 hours) in the conditioned medium from eIF4E^WT^ and eIF4E^KI^ primary melanoma cultures (WT-CM/KI-CM, n = 5 mice per group) or normal media as negative control (n = 5). One-way RM ANOVA, Tukey’s multiple comparisons test. **(E)** Inhibition of CD8^+^ T cells (% IFNγ production) by MDSCs at the indicated ratio in WT-CM or KI-CM (n = 6 per group), relative to corresponding MDSC-free control group. Two-way RM ANOVA with Sidak correction for multiple comparisons. **(F)** MDSC migration towards WT-CM, KI-CM (n = 5 per group) or normal media as negative control (n = 4). One-way ANOVA, Tukey’s multiple comparisons test. **(G)** MDSC migration towards media containing recombinant murine CCL5. One-way RM ANOVA, Tukey’s multiple comparisons test. **(H)** MDSC migration towards WT-CM or KI-CM (n = 5 per group) upon Maraviroc (100 nM) treatment. Two-way RM ANOVA with Sidak correction for multiple comparisons. **(I)** Percentage division of CD8^+^ T cells isolated from the ILNs of eIF4E^WT^ and eIF4E^KI^ tumor-bearing mice (n = 6 per group), cultured alone or with B16-F10 melanoma cells at indicated ratio. Two-way ANOVA with Sidak correction for multiple comparisons. One data point was excluded (Grubbs’ test). **(J)** Percent viability of B16-F10 cells co-cultured with T cells from eIF4E^WT^ and eIF4E^KI^ tumor-bearing animals (n = 4 per group). Two-way ANOVA with Sidak correction for multiple comparisons. See also Figure S5I. **(K)** Percent viability of B16-F10 cells, silenced or not for Melan-A (*siMlana-1*, see Table S4), co-cultured with T cells from eIF4E^WT^ (n = 4) and eIF4E^KI^ tumor-bearing animals (n = 5), relative to corresponding control groups. Two-way RM ANOVA with Sidak correction for multiple comparisons. For *ex vivo* assays **(C-K)**, all tumor-bearing mice were sacrificed between Day 35 and Day 38. All values are represented as mean ± SEM.

We hypothesized that the augmented CD8^+^ T cell influx and diminished presence of MDSCs in eIF4E^KI^ melanomas might be functionally associated with the anti-inflammatory signature identified in the conditioned media harvested from the melanomas lacking phospho-eIF4E (Figure 4B and S4A). To test this, we devised a number of *ex vivo* immune cell-based assays (Figure 5C). The ability of CD8^+^ T-cells to produce IFNγ was enhanced in the presence of eIF4E^WT^ tumor-conditioned media compared to normal culture media (Figure 5D), an effect that was significantly more robust when conditioned media obtained from the eIF4E^KI^ melanomas was used (Figure 5D). There was no differential effect on the proliferation of CD8^+^ T cells cultured in the conditioned media from eIF4E^WT^ or eIF4E^KI^ melanomas (Figure S5F). Moreover, tumor conditioned media also promoted the migration of MDSCs *ex vivo* (Figure 5F). However, the migration of MDSCs and their ability to suppress CD8^+^ T-cell function were both impaired under culturing conditions containing eIF4E^KI^ melanoma-derived conditioned media, compared to the eIF4E^WT^ melanoma conditioned media (Figure 5E, 5F and S5G). CCL5 is well known to promote the migration of MDSCs^69–72^. In our model, the addition of recombinant murine CCL5 to normal media promoted the migration of MDSCs, thus recapitulating the phenotype observed using eIF4E^WT^ tumor-derived conditioned media (Figure 5F and 5G). Strikingly, the CCR5 antagonist maraviroc (MVC)^73^, which blocks the CCL5-CCR5 axis, significantly decreased MDSC migration towards the eIF4E^WT^ conditioned media (Figure 5H). Such an effect was not observed using the eIF4E^KI^ conditioned media, which contains low CCL5 levels (Figure 5H). Together, these data suggest that the activation of the CCL5/CCR5 axis, regulated by phospho-eIF4E, is crucial for the recruitment and immune suppressive function of MDSCs.

Additional important factors contributing to the lack of T cells in melanomas have been proposed, such as a low presence of tumor antigens, including Melan-A and GP100 in highly dedifferentiated melanomas^30,31^. Therefore, we hypothesized that the increased presence of CD8^+^ T-cells and Granzyme B-positive cells in the eIF4E^KI^ melanomas may be a consequence of retention of melanocytic antigens in these tumors. To test this, we next isolated T cells from the sentinel (inguinal) lymph nodes of eIF4E^WT^ or eIF4E^KI^ tumor-bearing mice at Day 36 post 4HT administration, when sentinel lymph node metastasis is detectable in both mouse cohorts (Figure 5C, right panel). CD8^+^ T cells isolated from the eIF4E^KI^ lymph nodes were more proliferative compared to T cells isolated from the eIF4E^WT^ lymph nodes, when cultured alone or with B16-F10 melanoma cells that express high levels of Melan-A and GP100 (Figure 5I). Furthermore, using an *ex vivo* CD8^+^ cytotoxicity assay, we showed that B16-F10 melanoma cells were less viable when cultured in the presence of eIF4E^KI^ CD8^+^ T cells, compared to culturing them with WT CD8^+^ T cells (Figure 5J, S5H and S5I). The eIF4E^KI^ CD8^+^ T cells are highly functional against the B16-F10 melanoma cells, as increasing the T cell to tumor cell ratio lead to more robust tumor cell killing, while T cells isolated from the lymph nodes of non-tumor bearing mice showed minor tumor cell killing (Figure 5J and S5I). Finally, knockdown of Melan-A in B16-F10 melanoma cells significantly decreased the tumor cell killing ability of eIF4E^KI^ CD8^+^ T cells (Figure 5K, S5J and S5K). These data suggest that the retention of Melan-A expression in the eIF4E^KI^ melanomas leads to their recognition and eradication by CD8^+^ T cells.

### Both tumor cell-intrinsic and -extrinsic phospho-eIF4E are important for melanoma outgrowth and metastasis

In our eIF4E^KI^ autochthonous mouse model of melanoma, phospho-eIF4E is depleted in both the melanoma cells and the cells that comprise the TME. To discriminate between any potential tumor-intrinsic and -extrinsic effects of phospho-eIF4E on melanoma outgrowth, we used the MDMel melanoma-derived cells, which can be injected into immune competent C57BL/6 mice (Figure 6A). Strikingly, MDMel-KI-derived melanomas, where only the tumor cells are devoid of phospho-eIF4E, showed a significant defect in tumor outgrowth compared to MDMel-WT-derived melanomas (Figure 6B and 6C). The spleens of mice bearing MDMel-KI-derived melanomas were significantly smaller than those from the MDMel-WT tumor cohort (Figure S6A).

**Figure 6.**
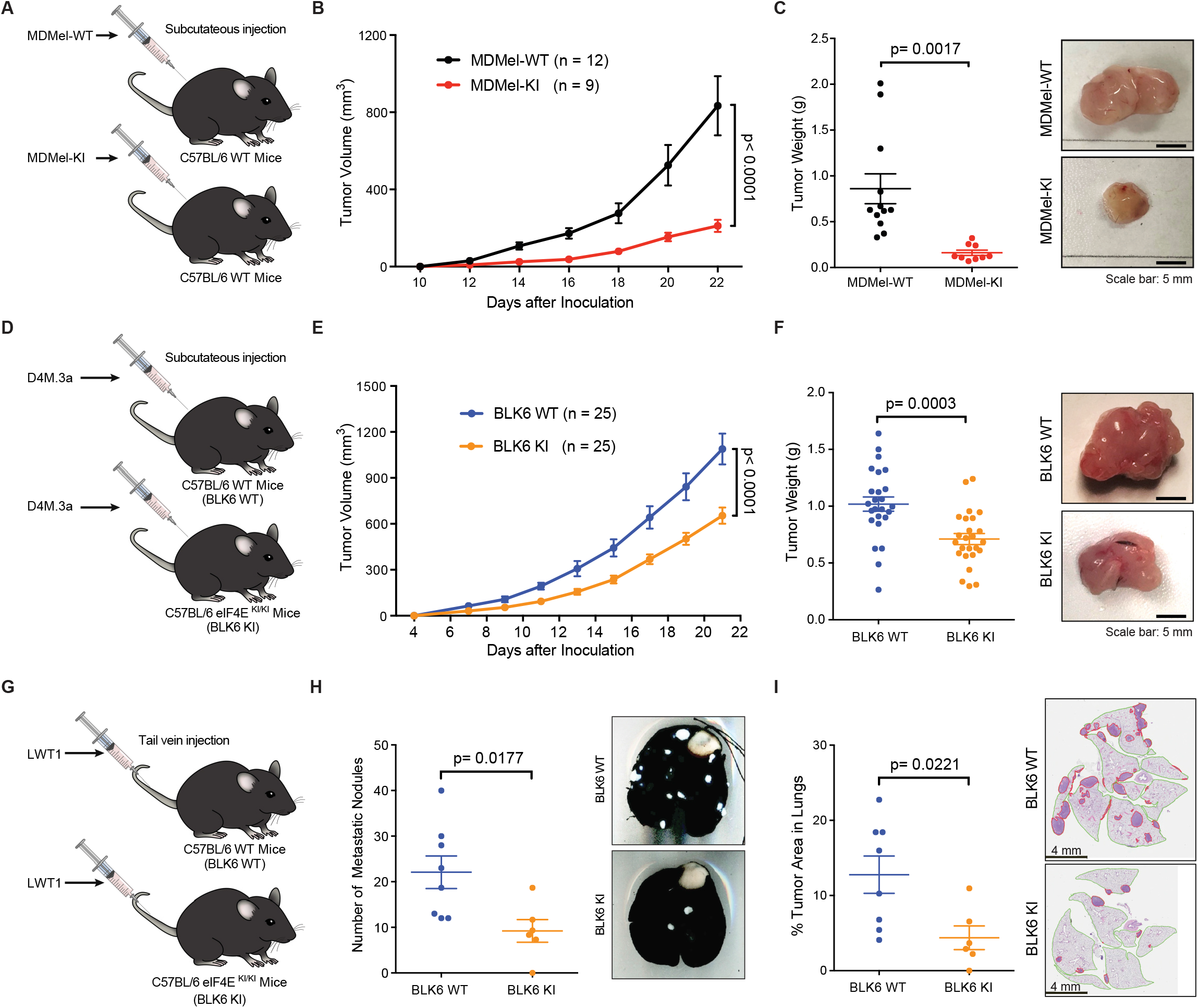
Tumor cell-intrinsic and -extrinsic phospho-eIF4E contributes to melanoma outgrowth and metastasis. **(A, D, G)** Schematic of the experimental design. **(B)** Growth curve of MDMel-WT (n = 12) and MDMel-KI (n = 9) derived melanomas. Two-way ANOVA. **(C)** Primary tumor weight (left) and representative pictures (right) of MDMel-WT (n = 12) and MDMel-KI (n = 9) derived melanomas (Day 22). Two-sided unpaired t test. **(E)** Growth curve of D4M.3a-derived melanomas in BLK6 WT and BLK6 KI mice (n = 25 per genotype). Two-way ANOVA. **(F)** Primary tumor weight (left) and representative pictures (right) of D4M.3a-derived melanomas in BLK6 WT and BLK6 KI mice (Day 21, n = 25 per genotype). Two-sided unpaired t test. **(H)** Number of metastatic nodules (left) and representative pictures of India Ink-inflated lungs (right) from BLK6 WT mice (n = 8) or BLK6 KI mice (n = 6) after tail vein injection of LWT1 cells (Day 21). Two-sided unpaired t test. **(I)** Percentage of tumor area (left) and representative images of H&E-stained lung sections (right) from BLK6 WT mice (n = 8) or BLK6 KI mice (n = 6) after tail vein injection of LWT1 cells (Day 21). Two-sided unpaired t test. All values are represented as mean ± SEM.

To dissect the relative contribution of phospho-eIF4E in the TME to melanoma outgrowth and metastasis, we injected D4M.3a melanoma cells subcutaneously into wild-type C57BL/6 (BLK6 WT) mice or phospho-eIF4E deficient C57BL/6 eIF4E^S209A/S209A^ (BLK6 KI) mice (Figure 6D). D4M.3a grew relatively slower in a host that lacks phospho-eIF4E, compared to grown in BLK6 WT mice (Figure 6E and 6F). No difference in spleen weight was found between BLK6 WT and BLK6 KI hosts (Figure S6B). Moreover, in a classic experimental model of lung metastasis^74–76^, when we injected BRAF^V600E^-driven murine melanoma cells LWT1 into the tail vein of BLK6 WT mice or BLK6 KI mice (Figure 6G), tumor burden was significantly decreased in the lungs of phospho-eIF4E deficient mice (Figure 6H and 6I).

Together, these data suggest that the phosphorylation of eIF4E in melanoma cells and in the cells that comprise the TME, both contribute to facilitating melanoma outgrowth and metastasis. Thus, the systemic inhibition of phospho-eIF4E, using a MNK1/2 inhibitor, could be therapeutically beneficial for patients with melanoma.

### Inhibition of MNK1/2-mediated eIF4E phosphorylation sensitizes melanoma to immune checkpoint blockade

Based on the data presented thus far, we propose that the protection of the eIF4E^KI^ mice against melanoma is likely a combined effect of phospho-eIF4E deficiency in the melanoma itself, but also in the cells of the TME. Inadequate tumor T cell influx, increased expression of T cell exhaustion markers, MDSC tumor infiltration, phenotype plasticity and dedifferentiation, can all underpin resistance to immunotherapies such as anti-PD-1^20,77–80^. Our data support the concept that blocking the phosphorylation of eIF4E reverses melanoma plasticity and increases CD8^+^ TILs. We thus suggest a promising strategy to block phospho-eIF4E systemically, using MNK1/2 inhibitors, to augment responses to immune checkpoint inhibitors (ICI). Concomitant with repressed phospho-eIF4E, the MNK1/2 inhibitor SEL201 significantly decreased the invasion and expression of NGFR and CCL5 in MDMel-WT melanoma cells (Figure 7A, 7B, S7A and S7B). Furthermore, SEL201 treatment of MDMel-WT cells faithfully recapitulates the profound changes in the cytokine/chemokine mRNA profile that we observed in the MDMel-KI cells (Figure 4D and S7A). This includes down-regulation of *Angpt2*, *Angptl4*, *Ccl2*, *Ccl12*, *Mmp9*, and unchanged *Ccl5* mRNA (Figure S7A). Given that the expression of *Angpt2* and *Angptl4* are not altered following *Ngfr* knockdown (Figure 4H), our results here indicate that these proteins are regulated by the MNK1/2-eIF4E axis, but are independent of NGFR. Notably, SEL201 treatment failed to alter the expression of *Igfbp2* and *Igfbp6* (Figure S7A).

**Figure 7.**
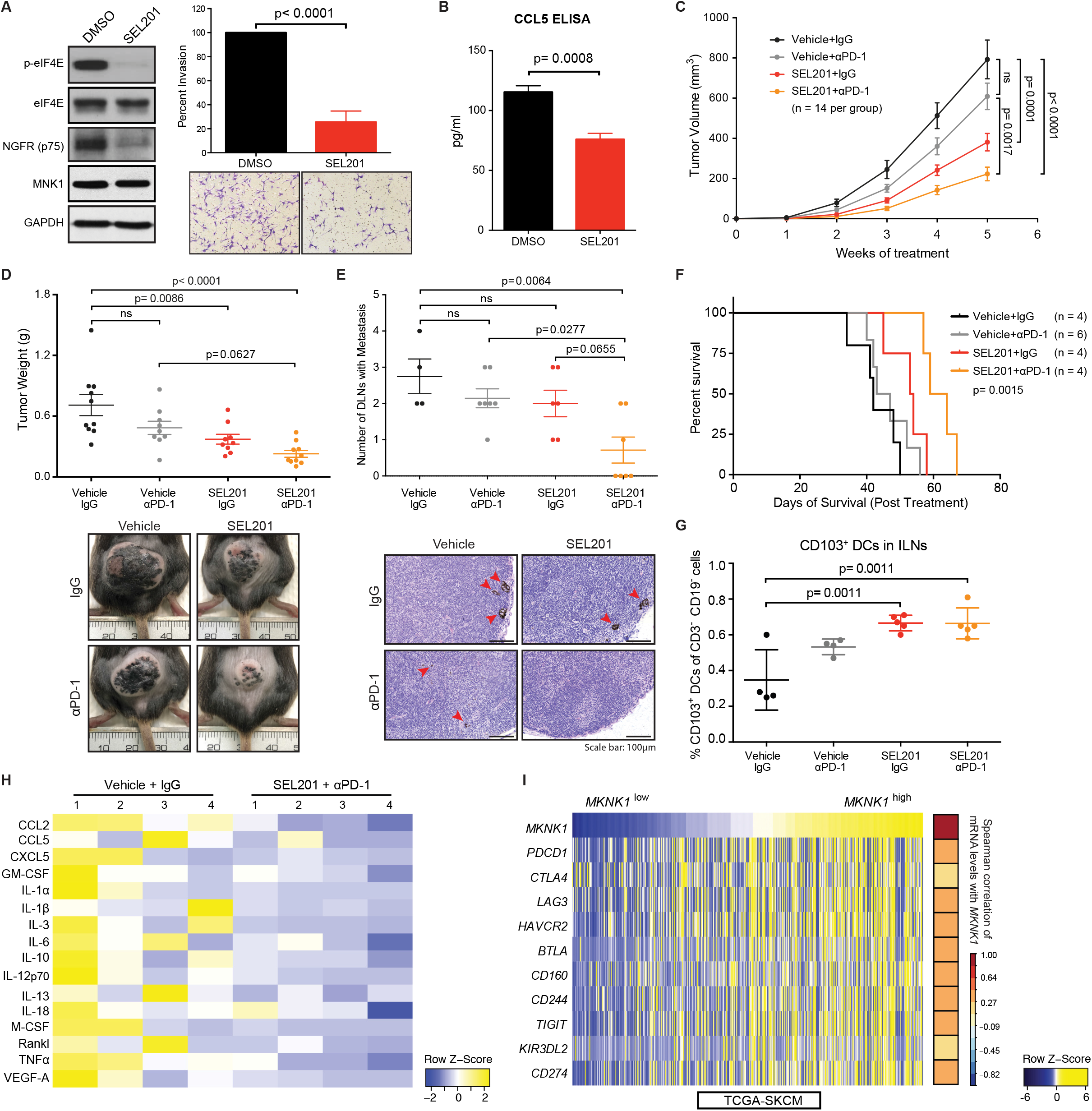
MNK1/2 inhibitor sensitizes melanoma to anti-PD-1 immunotherapy. **(A and B)** Characterization of melanoma cells treated with SEL201 (2.5 μM) *in vitro*. **(A)** Left: Western blot analysis of the indicated proteins in DMSO- or SEL201-treated MDMel-WT cells. Right: Percent invasion of SEL201-treated cells relative to control (top) and representative images with original magnification (×10, bottom). **(B)** CCL5 concentration in the conditioned medium of DMSO- or SEL201-treated MDMel-WT cells. Data are represented as mean ± SD (n=3). Two-sided unpaired t-test. **(C-H)** Evaluation of SEL201 treatment in combination with anti-PD-1 monoclonal antibody (αPD-1) *in vivo*. **(C)** Melanoma growth in mice administered control (Vehicle+IgG), monotherapies (Vehicle +αPD-1 or SEL201+IgG) and combination therapy (SEL201+αPD-1). Two-way RM ANOVA, Tukey’s multiple comparisons test. **(D)** Tumor weight (top) and representative pictures of melanomas (bottom) in each group (5-week treatment). Two-way ANOVA, Tukey’s multiple comparisons test. **(E)** Number of metastasis-positive CLNs per mouse (top) and representative images of H&E-stained CLNs (bottom) are presented for each group (5-week treatment). Two-way ANOVA, Tukey’s multiple comparisons test. **(F)** Kaplan–Meier curves showing overall survival of mice in each group. Log-rank test. **(G)** Percentage of CD103^+^ dendritic cells out of CD45^+^ CD3^−^ CD19-cells in the ILNs from animals in each indicated group (5-week treatment). Two-way ANOVA, Tukey’s multiple comparisons test. **(H)** Relative plasma cytokine and chemokine levels detected in the control (Vehicle+IgG) and combination therapy (SEL201+αPD-1) groups (treatment for 5 weeks). Plasma cytokine/chemokine concentration was detected using the MAGPIX multiplexing system. **(I)** Expression of *MKNK1* and immune exhaustion markers (HTSeq - FPKM) in GDC TCGA Melanoma dataset (SKCM, n = 472). Spearman rank-order. For *in vivo* experiments **(C-H)**, all values represent the mean ± SEM, with numbers of animals in each group (minimum n = 4) indicated in the figure.

Next, we combined SEL201 treatment with anti-PD-1 immunotherapy *in vivo*. *BRaf^CA/+^/Pten^lox/lox^* (eIF4E^WT^) melanomas were initiated, and when the melanomas were first visible the mice were administered SEL201 alone or in combination with an anti-PD-1 monoclonal antibody. Anti-PD-1 monotherapy alone did not significantly affect macroscopic melanoma outgrowth, metastasis, and the survival of animals (Figure 7C-7F and S7C-S7F), consistent with prior studies using this mouse model^68,81^. In contrast, SEL201 reduced both the primary melanoma outgrowth and distant lymph node metastasis (Figure 7C-7E and S7D), improved the survival of animals (Figure 7F) and increased antigen-presenting CD103^+^ dendritic cells in the sentinel lymph nodes (Figure 7G). SEL201 administration in mice showed robust circulating levels of drug (Figure S7C) and on-target engagement, as shown by the repression of phospho-eIF4E expression in the melanomas (Figure S7D). These data recapitulate our observations in the phospho-eIF4E deficient *BRaf^CA/+^/Pten^lox/lox^* (eIF4E^KI^) mice, and further supports that systemic inhibition of phospho-eIF4E by blocking MNK1/2, has efficacy in modulating both the tumor and TME in pre-clinical cancer models^48,82–85^. Strikingly, the combination therapy of SEL201 and anti-PD-1 monoclonal antibody significantly reduced primary melanoma growth, local and distant lymph node metastasis, and increased the overall survival of tumor-bearing *BRaf^CA/+^/Pten^lox/lox^* mice compared to either of the therapies alone, without causing any overt toxicity (Figure 7C-7F and S7C-S7E).

Analysis of plasma cytokine and chemokine levels revealed that the robust anti-tumor effect of SEL201 combined with anti-PD-1 therapy was associated with enhanced peripheral anti-tumor immunity, characterized by a decrease in the immunosuppressive cytokine/chemokine signature, including IL-6, CCL2 and CCL5 (Figure 7H). Such a peripheral cytokine/chemokine signature might have prognostic value to predict responders of combination therapy with MNK1/2 inhibitors and immune checkpoint blockade. Strikingly, although demonstrating a better anti-tumor response, the combination therapy led to a profound decrease in a subset of circulating cytokines that have been previously associated with immune-related adverse events due to immune checkpoint blockade (Figure S7G)^86,87^. Furthermore, in contrast to the *BRaf^CA/+^/Pten^lox/lox^* eIF4E^WT^ mice, the *BRaf^CA/+^/Pten^lox/lox^* eIF4E^KI^ mice are sensitive to anti-PD-1 monotherapy, showing a marked inhibition in macroscopic tumor outgrowth (Figure S7H).

Exhausted T cells can be defined by their expression of immune checkpoints, such as PD-1, LAG3, TIGIT^19,20^. We used the TCGA SKCM cohort to interrogate whether *MKNK1* expression in patient-derived melanomas correlates with markers of immune exhaustion. Strikingly, high *MKNK1* expression was positively correlated with an immune escape gene signature, comprised of *PDCD1*, *CTLA4*, *LAG3*, *HAVCR2*, *BTLA*, *CD160*, *CD244*, *TIGIT*, *KIR3DL2* and *CD274* (Figure 7I). Curiously, melanomas with elevated *MKNK1* expression also correlated with high expression of a panel of genes reflecting MDSC infiltration (*ITGAM*, *CD33*, *S100A9*, *CCR5*, *CD40*, *IL10*, *IL4R*, *IL1B*, *IL10*, *CSF2*, *CD14* and *FUT4*) (Figure S7I)^63–66^. These results suggest that MNK1 is part of a regulatory network to drive immune suppression in human melanoma, and support ongoing clinical trial work wherein MNK1/2 inhibitors are being tested in combination with immune checkpoint blockade (NCT03616834).

## Discussion

We have shown an integral role for the MNK/12-eIF4E axis in regulating melanoma plasticity. Abrogation of eIF4E phosphorylation attenuated melanoma phenotype switching via a mechanism involving the translational repression of NGFR. Inhibition of phospho-eIF4E in cutaneous melanoma led to increased/preserved expression of melanocytic antigens Melan-A and GP100, decreased production of pro-inflammatory and immunosuppressive cytokine/chemokines and reduced tumor infiltration of MDSCs. Those changes, in turn, resulted in the observed diminished melanoma outgrowth, reduced invasiveness and metastasis, and increased sensitivity to anti-PD-1 immunotherapy, which all resulted from repressed MNK1/2-eIF4E axis activity.

Our findings are consistent with prior work demonstrating that BRAF and/or MEK inhibitors can synergize with immune checkpoint blockade in melanoma, via increased antigen expression, decreased pro-inflammatory cytokine/chemokine production and improved T cell infiltration^79,88–93^. However, such combinations can result in a high rate (~73%) of grade 3/4 treatment-related adverse events in patients with melanoma^92^. As MNK1/2 inhibitors have reached clinical trials (NCT03616834, NCT03690141, NCT04261218), it will be important to determine whether targeting MNK1/2, downstream of activated BRAF/MEK/ERK, is potentially better tolerated. Encouragingly, several peripheral cytokines recently shown to predict immune-related toxicity of immune checkpoint blockade^86,87^ were repressed when *BRaf^CA/+^/Pten^lox/lox^* mice were treated with combined anti-PD-1 and SEL201. Those studies coupled with recent work reporting that targeting phospho-eIF4E and the eIF4F complex can reduce the expression of PD-L1^82,94^, highlight the need to further understand the role of translational control in melanoma.

EMT plays an important role in tumor immune escape^95–97^. In this context, translational control, including the MNK1/2-eIF4E axis, has been implicated in promoting an EMT across different tumor types^10,85,98,99^. Our study links the MNK1/2-eIF4E axis with immune escape of melanoma though an EMT-like phenotype switch. Given the importance of the melanocyte differentiation antigens (e.g. Melan-A) in melanoma immunogenicity and in favoring a good response to immune checkpoint inhibitors^28,31,37,100–104^, the relevance of combined blockade of phenotype switching and immune checkpoints in melanoma should not be neglected. For instance, increased expression of these melanocytic antigens stimulate a vigorous T cell response in melanoma and CD8^+^ T cells specific for these antigens effectively induce regression of melanoma metastases in patients^105–107^. Accordingly, the dedifferentiation of melanoma has been demonstrated as a major mechanism for the acquired resistance of melanomas to immunotherapies, including adoptive T cell transfer^28,30,31,104^. Indeed, our study shows that blocking the phosphorylation of eIF4E can resist the dedifferentiation of melanoma, leading to increased expression of Melan-A (and GP100), which significantly enhanced the infiltration of T cells and their functions in tumor recognition and elimination.

Increased production of pro-inflammatory cytokine/chemokines and infiltration of T cell inhibitory MDSCs are important characteristics of the immunosuppressed TME of dedifferentiated melanomas^38,61^. We were able to identify a unique subset of cytokines/chemokines (ANGPT2, CCL2, CCL5, CCL12, IGFBP-2, IGFBP-6, IL-6, MMP-9) whose expression was tightly regulated by the phosphorylation of eIF4E (Figure 4A, 4B and S4A). Among these genes, few have been closely investigated in the context of melanoma phenotype switching. Strikingly, a re-analysis of previously published RNAseq data, derived from melanomas with an invasive phenotype^53–55,61,62^, highlighted that the phospho-eIF4E-regulated cytokines/chemokines are also phenotype switch-related targets^53–55,61,62^. Among these factors, we report CCL5, a chemokine that plays a crucial role in MDSC recruitment, as a novel translationally controlled mRNA by the phosphorylation of eIF4E. Similarly, other cytokines, including CCL2, CCL12, IL-6 and MMP-9, were all decreased in the eIF4E-deficient melanomas, which together led to a less immunosuppressive TME (Figure 4 and 5). Interestingly, two unexpected families of secreted factors appear to be regulated under specific conditions of repressed phospho-eIF4E, the angiopoietins/angiopoietin-like proteins and the IGFBPs. The expression of angiopoietins was down-regulated following the MNK1/2 inhibitor SEL201 treatment, but not siRNA-mediated knockdown of *Ngfr*, while IGFBPs responded to *Ngfr* knockdown (48 hours), but not to a 24-hour SEL201 treatment (Figure 4D, 4H and S7A). Notably, we recently reported that ANGPTL4, which was not originally on the cytokine array used in this study, was regulated via MNK1^108^. Our results here further suggest that *Angptl4*, as well as *Angpt2* (potentially, *Angpt1* and *Angptl3*), are regulated downstream of MNK1/2-eIF4E signaling, but independent of NGFR. The expression of *ANGPT2* and *ANGPTL4* are positively correlated with *NGFR* expression (Figure 4I), indicating that they might be regulated through the same upstream signaling. The significance of the robust repression of IGFBP-2 and IGFBP-6 expression detected in the conditioned media from phospho-eIF4E deficient melanomas (Figure 4B and S4A) warrants our further investigation.

Mechanistically, both tumor-intrinsic and TME-mediated regulation of phenotype switching has been proposed^28–31,33,36–38,109^. We identified NGFR, a receptor that links intracellular signaling with extracellular factors in TME, as a key downstream translational target of phospho-eIF4E that drives phenotype switching. Our results show that the expression of the NGFR protein is downregulated via genetic and pharmacologic blockade of phospho-eIF4E. Short-term inhibition of phospho-eIF4E using the MNK1/2 inhibitor SEL201 repressed NGFR protein, but not mRNA expression (Figure 7A and S7A), indicating that the changes in *Ngfr* mRNA expression observed in the context of genetic repression of phospho-eIF4E may involve a mechanism that takes place at a later time point (Figure 7A and S7A). Our interrogation of the TCGA melanoma data set led us to remark that *NGFR* gene expression correlates not only with the dedifferentiation gene signature, but also with the phospho-eIF4E-regulated cytokine/chemokine signature that we reported in our mouse model, and with a set of genes associated with MDSC infiltration (Figure 4I). Notably, no significant correlation between NGFR and CCL5 gene expression was observed. This is to be expected, as we reported that the expression CCL5 is regulated by phospho-eIF4E at the mRNA translation level. Furthermore, CCR5, the CCL5 receptor, also shows a strong correlation with NGFR expression, providing additional evidence that the CCL5-CCR5 axis might play an important role in melanoma immune escape.

In this study, we showed that the phosphorylation of eIF4E has a critical role in phenotype switching by controlling the translation of NGFR mRNA, which encodes a key mediator of the invasive phenotype switch. Intriguingly, the expression of phospho-eIF4E inversely correlated with a loss of Melan-A expression in patient-derived samples, demonstrating a previously unknown association between the activation of the MNK1/2-eIF4E axis and the dedifferentiation (phenotype switching) of human melanoma. Finally, soluble inflammatory factors released by primary melanomas are controlled by phospho-eIF4E to help shape an immunosuppressive microenvironment. Specifically, we uncover a mechanism of regulation of immune suppression mediated by MDSCs by translational control of *Ccl5* mRNA in melanoma. In toto, our results here provide evidence that, although blocking phospho-eIF4E alone shows efficacy in our preclinical melanoma model, these agents also serve as potent immune modulators that will likely be most therapeutically beneficial when combined with immune checkpoint blockade.

## Methods

### Mice

C57BL/6N mice and NOD/SCID mice (strain code 394) were purchased from Charles River Laboratories. C57BL/6 *Tyr::CreER*/*BRaf*^CA/+^/*Pten*^lox/lox^ mice (referred to as eIF4E^WT^ mice in this paper)^39^ was a gift from David Dankort. C57BL/6 eIF4E^S209A/S209A^ mice^9^ was a gift from Nahum Sonenberg. The *Braf^CA/+^/Pten^lox/lox^/eif4e^S209A/S209A^* mice (referred to as eIF4E^KI^ mice in this paper) were generated by crossing the C57BL/6 eIF4E^S209A/S209A^ mice and the C57BL/6 *Tyr::CreER*/*BRaf*^CA/+^/*Pten*^lox/lox^ (eIF4E^WT^) mice. Genotyping was performed with indicated primer sets (See Table S2).

For tumor induction, the eIF4E^WT^ and eIF4E^KI^ mice (6-8 weeks old) were treated with 2 μl 4-HT (Sigma) at 5 mM working concentration for 3 continuous days. Tumors were subsequently measured in length (L), Width (W) and Thickness (H) (mm). Tumor volumes (V) were calculated based on the formula V=3.1416/6*L*W*H. Inguinal lymph node sizes were measured by ultrasound and analyzed and calculated using the software Vevo. For subcutaneous injections, tumor cells in PBS were injected to the right flank of the mice. Tumors were measured in length (L) and Width (W) (mm). Tumor volumes (V) were calculated based on the formula V=3.1416/6*L*W^2^. For metastasis experiments, 1 million LWT1 cells were injected through tail veins of C57BL6 mice. For combination treatment, eIF4E^WT^ mice were randomly grouped before 4-HT administration. 12-15 days after 4-HT induction, when pigmented lesions started to develop, mice were treated with SEL201 and/or anti-mouse PD-1 monoclonal antibody (RMP1-14), with vehicle and/or IgG isotype as controls, repectively. SEL201 was dissolved in DMSO and then diluted in Captisol (Ligand) for administration by oral gavage at 75 mg/kg bodyweight per mouse per day, 5 days per week. The anti-mouse PD-1 monoclonal antibody and IgG isotype control were diluted in PBS and administrated through intraperitoneal injection at 10 mg/kg bodyweight per mouse per day, once per week. Animals were euthanized at indicated time points. Tumors and other organs were resected, processed right away or stored either in 10% formalin or at −80°C for indicated use. Lungs were inflated with either 10% formalin for storage or with India ink for visualization of lung metastasis. All mice used in this study are males. Animal experiments were conducted following protocols approved by McGill University Animal Care and Use Committee.

### Cells and Reagents

The <u>M</u>ontreal-<u>D</u>erived murine <u>Mel</u>anoma (MDMel) cell lines, MDMel-WT^88^, MDMel-WT^73^, MDMel-KI^61^ and MDMel-KI^58^ were generated similarly as murine melanoma cell line D4M.3a used in this paper^46^. Briefly, primary tumors from eIF4E^WT^ and eIF4E^KI^ mice were resected, minced and digested with collagenase A. Cells were cultured in Advanced DMEM/F12 media supplemented with 5% FBS, 1x GlutaMax and 1x Antibiotic-Antimycotic (Penicillin/Streptomycin/Amphotericin B cocktail, ThermoFisher) for 48 hours before subsequently trypsinized and injected to SCID/Beige mice subcutaneously. The newly formed tumors were again resected, minced and digested with collagenase A. Cells were cultured in Advanced DMEM/F12 media supplemented with 1% FBS, 1x GlutaMax and 1x PSA until they were tested as PTEN negative. All MDMel cell lines were cultured in Advanced DMEM/F12 media supplemented with 5% FBS, 1x GlutaMax and 1x Pen/Strep. Unless otherwise specified, All MDMel-WT cells refer to the MDMel-WT^73^ cell line; All MDMel-KI cells refer to the MDMel-KI^61^ cell line. The D4M.3a cell line was a kind gift from Dr. Brinckerhoff, C. E., and was cultured in Advanced DMEM/F12 media supplemented with 5% FBS, 1x GlutaMax and 1x Pen/Strep. The LWT1 cell line was a kind gift from Dr. Mark J. Smyth, and was cultured in RPMI supplemented with 10% FBS, 1x GlutaMax and 1x Pen/Strep. The B16-F10 cell lines, was a kind gift from Dr. Michael Pollak (Lady Davis Insitute). The human HBL-*shCtrl* and HBL-*shMKNK1/2* cell lines were generated as previously described^48^.

### Western Blotting

Frozen tumor samples were ground in liquid nitrogen. Ground tissue or cells were lysed with RIPA buffer (150 mmol/L Tris-HCl, pH 7, 150 mmol/L NaCl, 1% NP-40, 1% sodium deoxycholate, 0.1% SDS) supplemented with protease and phosphatase inhibitors (Roche) as previously described. Equal amounts of protein samples were loaded, separated on 10% SDS-PAGE gels, transfered to PVDF membranes and probed with corresponding antibodies. The anti-MITF monoclonal C5 antibody^110^ was a kind gift from Dr. David E Fisher. Detailed antibody information is listed in Table S1.

### Polysome profiling

Polysome profiling was performed as previously described^48^. Briefly, MDMel-WT^73^, MDMel-KI^61^, WT-MEF and KI-MEF cells were serum starved over night and then serum stimulated for 2 hours. Cells were treated with cycloheximide (100 μg/ml) 5 minutes before harvesting, then washed in cold PBS containing 100 μg/ml cycloheximide, followed by centrifugation for 5 minutes at 390 g. Cell pellets were lysed in hypotonic buffer: 5 mM Tris-HCl (pH 7.5), 2.5 mM MgCl_2_, 1.5 mM KCl, and 1x protease inhibitor cocktail, containing 1 mM DTT and RNase inhibitor (100 U). Samples were kept on ice for 12 minutes, then centrifuged at 13,523 g for 7 minutes. The supernatants were loaded onto 10%–50% sucrose density gradient and centrifuged at 260,110 g for 2 hours at 4°C. The polysomal fractions were monitored and collected using a Foxy JR ISCO collector.

### Quantitative real-time PCR

For polysomal fractions, RNA was extracted from fraction 7 to 17 as well as input samples using the TRIzol method. For cell pellets, RNA was prepared using E.Z.N.A. total RNA isolation kit (OMEGA Bio-Tek). cDNA was prepared from 1 mg of total RNA, using iScript cDNA Synthesis Kit (Bio-Rad). Target genes were quantified using the Applied Biosystems 7500 Fast Real-Time PCR System with SYBR Green. Primers used for qPCR are listed in Table S3.

### PCR

cDNAs converted from RNAs of each polysomal fractions were used as templates. PCR was performed using the AccuStart II GelTrack PCR SuperMix kit (QuantaBio) and allowed to run 35 amplification cycles. Amplified cDNA fragments of indicated targets were analyzed by electrophoresis on a 1% denaturing agarose gel. Primers used for PCR are listed in Table S2.

### RNA interference

siRNAs were transfected into MDMel-WT or B16-F10 cells using transfection reagent Lipofectamine RNAiMAX following the manufacturer’s instructions. All siRNA sequences are listed in Table S4.

### Invasion Assay

MDMel-WT^88^, MDMel-WT^73^, MDMel-KI^61^ and MDMel-KI^58^ cells were serum starved overnight. Transwells were coated with Collagen I at 20 μg/ml for 30 mins. 30,000 cells in serum-free media were seeded into each transwell and were allowed to migrate and invade to the bottom chamber with media containing 5% FBS for 24 hrs. Cells were then fixed with 5% glutaraldehyde (Sigma) and stained with 0.5% crystal violet. Non-migrated cells were manually removed. The remaining migrated cells were quantified manually.

### Gelatin zymography assay

MMP9 levels in cell culture supernatant were detected by gelatin zymography as previously described^108^. Briefly, a 10 ml cell culture supernatant was concentrated to 0.2 ml by using Amicon Ultra-15 centrifugal spinning units (Millipore). Concentrated supernatants were subsequently separated with 7.5% acrylamide gels containing 0.1% gelatin A (Thermo Fisher) and digested bands by MMP9 were revealed with 0.25% Coomassie blue staining.

### Immunohistochemistry and scoring

Human melanoma tissue samples were acquired in collaboration with G. Ghanem (Institut Jules Bordet), and written informed consent from all patients was obtained in accordance wih the Declaration of Helsinki. Immunohistochemistry staining of human patient samples were performed on a Ventana Discovery Benchmark XT while mouse samples were performed as previously described^48^. Briefly, formalin-fixed, paraffin-embedded tumor sections were stained with indicated antibodies (see Table S1) at 1:50 dilution, followed by a standard Fast Red detection protocol. Hematoxylin-counterstained slides were mounted with coverslips. Staining intensity was determined by a clinically certified pathologists. All pathologists were blinded to all clinical data and antibodies used for IHC.

### Tumor conditioned media

eFI4E^WT^ and eIF4E^KI^ melanomas were resected at Day 35-Day 38 after 4-HT administration. Tumors were minced and digested in collagenase A to obtain single cell suspension and cultured in advanced DMEM/F12 supplemented with 5% FBS, 1x glutaMax and 1x Antibiotic-Antimycotic at ~3-4 million cells per 15 cm dish overnight. Then cells were washed once with PBS and changed to serum free DMEM/F12 media supplemented with 1x Antibiotic-Antimycotic for 24 hours. Media were then harvested and protein concentrations were adjusted based on Bradford measurement. These media were then used for cytokine array, ELISA, or supplemented with 10% FBS to make tumor conditioned media for *ex vivo* immune cell assays.

### Cytokine Array and ELISA

Cytokine array was performed with Proteome Profiler Mouse XL Cytokine Array kit (R&D) following the manufacturer’s protocols. Mouse CCL5 and IL-6 ELISA were performed with the Mouse/Rat CCL5/RANTES Quantikine ELISA Kit (R&D) and the Mouse IL-6 Quantikine ELISA Kit (R&D), respectively, following the manufacturer’s protocols.

### Immunophenotyping

Tumors were minced and digested in 4 mg/ml collagenase A (Sigma Aldrich) at 37°C for 1 hour to obtain single cell suspension and were stained with indicated antibodies (see Table S5) as well as a live/dead discrimination dye. Data were subsequently acquired at the Fortessa flow cytometer (BD Biosciences).

### T cell activation

T cells were isolated from spleens of naïve male C57BL/6 mice using the EasySep™ Mouse T Cell Isolation Kit (STEMCELL) and stained with 1.4 ng/ml carboxyfluorescein succinimidyl ester (CFSE). Cells were plated in CD3-coated 96-well plates (10 μg/ml CD3 in PBS for 30 min at 37 °C) at 100,000 cells/well and were allowed to expand for 72h in ImmunoCult™-XF T Cell Expansion Medium (STEMCELL) supplemented with 20 ng/μl IL-2 and 30% of eFI4E^WT^ or eIF4E^KI^ conditioned media. BD GolgiStop™ (Monensin) and BD GolgiPlug™ (Brefeldin A) were added at the manufacturer recommended concentrations 3-4 hours before harvest. Cells were stained with indicated antibodies (see Table S5) as well as a live/dead discrimination dye. Data were subsequently acquired at the Fortessa flow cytometer.

### MDSC/T cell co-culture assay

MDSC were isolated from spleens of tumor-bearing eFI4E^WT^ mice using the EasySep™ Mouse MDSC (CD11b^+^Gr1^+^) Isolation Kit (STEMCELL). MDSCs were seeded together with 75,000 T cells at indicated ratios in CD3-coated 96-well plates. Media used in this assay is indicated above (see T cell activation).

### MDSC migration assay

100,000 freshly isolated MDSCs were seeded in the Corning Transwell cell culture inserts in DMEM/F12 media supplemented with 10% FBS and 1x Antibiotic-Antimycotic. eFI4E^WT^ or eIF4E^KI^ conditioned media were added to the bottom chamber. Cells were cultured for 6 hours. Cells migrated in bottom chambers were manually counted.

### T cell/Tumor cell co-culture assay

B16/F10 melanoma cells were seeded in 24 well plates at 2,000 cells/well, or in 96 well plates at 400 cells/well one day before T cell extraction. T cells were extracted from inguinal LNs of eFI4E^WT^ or eIF4E^KI^ tumor bearing animals and were purified using the EasySep™ Mouse T Cell Isolation Kit (STEMCELL) and stained with 1.4 ng/ml CFSE. T cells were added to the tumor cells in ImmunoCult™-XF T Cell Expansion Medium (STEMCELL) supplemented with 20 ng/μl IL2 at indicated T cell to tumor cell ratios. Cells were then cocultured for another 72hours. T cells in 24 well plates were harvested to access proliferation and IFNγ production. T cells and dead tumor cells in 96 well plates were washed off by PBS and tumor cells were fixed with 4% PFA and stained with crystal violet. Percentage survival was accessed by OD value compared to tumor cells cultured alone.

### Statistical Analysis

*In vitro* data were represented as mean ± SD. *In vivo* and *ex vivo* data were represented as mean ± SEM. Prism software (GraphPad) was used to determine statistical significance of differences. The GDC TCGA Melanoma (SKCM) data was accessed through the UCSC Xena platform (Xenabrowser)^111^. Heat maps were generated through Heatmapper^112^. Two-sided Student’s t-test, one-way ANOVA or two-way ANOVA is used, as appropriate. Figure legends specify the statistical analysis used and define error bars. P values are indicated in the figures and P values < 0.05 were considered significant.

## Author Contributions

Designing research studies: F. Huang, S. V. del Rincón

Conducting experiments: F. Huang, C. Gonçalves, M. Bartish

Acquisition of data: F. Huang, C. Gonçalves, M. Bartish, J. Rémy-Sarrazin, Q. Guo, A. Emond, W. Yang, D. Plourde, J. Su, Y. Zhan, M. Attias, A. Galán, M. Mazan, M. Masiejczyk, J. Faber, E. Khoury, A. Benoit, N. Gagnon

Analysis and interpretation of data: F. Huang, C. Gonçalves, M. Bartish, A. Emond, M. G. Gimeno, M. Attias, A. Galán, T. Rzymski

Writing, review, and/or revision of the manuscript: F. Huang, M. Bartish, H. U. Saragovi, N. Sonenberg, I. Topisirovic, W. H. Miller, S. V. del Rincón.

Administrative, technical, or material support: C. Gonçalves, D. Dankort, C. A. Piccirillo, F. Journe, G. Ghanem

Study supervision: W. H. Miller and S. V. del Rincón.

## Acknowledgements

This research is funded by the Canadian Institutes for Health Research (grant PJT-162260 to SVDR and grants MOP-142281 and PJT-156269 to WHM and SVDR) and the Canadian Cancer Society (grant 703811 WHM). This work was also supported by the Terry Fox Research Institute-Montreal Cancer Consortium (TFRI – Grant #1084). The research was further supported by the Rossy Cancer Network. Development of MNK1/2 inhibitors by Ryvu Therapeutics has been co-financed by the National Centre for Research and Development, INNOTECH Program (INNOTECH-K1/HI1/I6/157438/NCBR/12). FH was endowed by MICRTP graduate studentships. QG was financed by a Cole Foundation Ph.D. fellowship, McGill Integrated Cancer Research Training Program (MICRTP) graduate studentship and a McGill Faculty of Medicine graduate studentship. AB was financed by a Cole Foundation Ph.D. fellowship. WY was sponsored by MICRTP graduate studentships. We thank Dr. David Fisher (Harvard Medical School, Boston, Massachusetts.) for his generous donation of the MITF antibody. We thank Christian Young, Naciba Benlimame, Lilian Canetti, Valeria Narykina, Sathyen A. Prabhu and Sylvain Roux for experimental advice and technical supports.

