## Supplementary material for "The MNK1/2-eIF4E axis drives melanoma plasticity, progression, and resistance to immunotherapy": Supp. Tables S1-S4

**Table S1. Detailed information of primary antibodies used for western blotting, immunohistochemistry and immunofluorescence staining.**

| Target | Antibody full name | Source | Experiment |
|---|---|---|---|
| Ki67 | Ki-67 (D2H10) Rabbit mAb (IHC Specific) | Cell Signaling | IHC staining |
| S100 | S100 Protein Ab-2 Rabbit Polyclonal Antibody | ThermoFisher | IHC staining |
| MITF | Anti-MITF monoclonal C5 antibody | Dr. David E Fisher Lab | Western blot |
| Phospho-eIF4E | Phospho-eIF4E (Ser209) Antibody | Cell Signaling | Western blot; IHC staining |
| eIF4E | Purified Mouse Anti-eIF-4E Clone 87/eIF-4E (RUO) | BD Biosciences | Western blot |
| Melan-A | Recombinant Anti-MelanA antibody [EPR20380] | Abcam | Western blot; IHC staining |
| GP100 | Rabbit Anti-Melanoma gp100 antibody [EP4863(2)] | Abcam | Western blot; IHC staining |
| Tyrosinase | Tyrosinase Monoclonal Antibody (T311) | Invitrogen | Western blot |
| Tyrp-1 | Recombinant Anti-TRP1 antibody [EPR13063] | Abcam | Western blot |
| Tyrp-2 | Recombinant Anti-TRP2/DCT antibody [EPR21986] | Abcam | Western blot |
| MNK1 | Mnk1 (C4C1) Rabbit mAb | Cell Signaling | Western blot |
| PTEN | PTEN (138G6) Rabbit mAb | Cell Signaling | Western blot |
| NGFR | Anti-p75$^{NTR}$ Antibody, ICD | Sigma-Aldrich | Western blot; IF staining |
| CD8 | CD8a Monoclonal Antibody (4SM15) | eBioscience | IF staining |
| Granzyme B | Rabbit Anti-Granzyme B antibody | Abcam | IHC staining |
| GAPDH | GAPDH (14C10) Rabbit mAb | Cell Signaling | Western blot |
| β-Actin | Monoclonal Anti-β-Actin (AC-15) | Sigma-Aldrich | Western blot |

### Table S2. Genotyping/PCR primers.

| Allele/Gene | Primer | Sequence (5'-3') | Amplicon (bp) |
|---|---|---|---|
| ***Tyr::CreER$^{T2}$*** | Fwd | CTGGAAGGGATTTTTGAAGCAAC | Mutant: ~375 |
| ***(TCE)*** | Rev | CCAACAAGGCACTGACCATCTGGTC | WT: No band |
| ***BRaf$^{CA}$*** | Fwd | GGAAAGCCTGTCACGGGTC | Mutant: 413 |
| | Rev | AGATTCGTATGTCCTCTGAAAGTC | WT: ~357 |
| ***Pten$^{flox}$*** | Fwd 1 | AAAAGTTCCCCTGCTGATGATTTGT | Mutant: ~400 |
| | Fwd 2 | CCCCCAAGTCAATTGTTAGGTCTGT | WT: ~342 |
| | Rev | TGTTTTTGACCAATTAAAGTAGGCTGTG | |
| ***Eif4e$^{KI}$*** | Fwd | TTTGAAATTGGTTTGTAAAGTTGG | Mutant (KI): ~420 |
| | Rev | GCAATGCAAGTCGAAATGTG | WT: ~313 |
| ***Ccl5*** | Fwd | GCCCACGTCAAGGAGTATTT | ~311 |
| ***NM_013653.3*** | Rev | CTGATTTCTTGGGTTTGCTGTG | |
| ***Ngfr*** | Fwd | GTCCACACTCCTTCTCTTACAC | ~509 |
| ***NM_033217.3*** | Rev | TCTCTCTCTCTCTCTCTCTCTCT | |
| ***Rplp0 (m36B4)*** | Fwd | TCATCCAGCAGGTGTTTGACA | ~151 |
| ***NM_007475.5*** | Rev | GGCACCGAGGCAACAGTT | |
| ***Actb*** | Fwd | GGCTGTATTCCCCTCCATCG | ~154 |
| ***XM_030254057.1*** | Rev | CCAGTTGGTAACAATGCCATGT | |

### Table S3. RT-qPCR primers.

| Gene name | Primer | Sequence (5’-3’) |
|---|---|---|
| ***Angpt2*** | Fwd | GATCTTCCTCCAGCCCCTAC |
| ***NM_007426.4*** | Rev | TTTGTGCTGCTGTCTGGTTC |
| ***Angptl4*** | Fwd | CTGAATATCACTTCTCGCCTACC |
| ***NM_020581.2*** | Rev | CCTGTCTCCAGTCAGTCAATATG |
| ***Ccl2*** | Fwd | TTTTGTCACCAAGCTCAAGAGA |
| ***NM_011333.3*** | Rev | ATTAAGGCATCACAGTCCGAGT |
| ***Ccl5*** | Fwd | CTGCTGCTTTGCCTACCTCT |
| ***NM_013653.3*** | Rev | CGAGTGACAAACACGACTGC |
| ***Ccl12*** | Fwd | AATCACAAGCAGCCAGTGTCC |
| ***NM_011331.3*** | Rev | TCAGCACAGATCTCCTTATCCAGT |
| ***Il6 NM_031168.2*** | Fwd | TGATGCACTTGCAGAAAACA |
| ***NM_001314054.1*** | Rev | ACCAGAGGAAATTTTCAATAGGC |
| ***Igfbp2 NM_001310659.1*** | Fwd | CGAGTGCCATCTCTTCTACAAC |
| ***NM_008342.3*** | Rev | GAGCTCAGTGTTGGTCTCTTTC |
| ***Igfbp6*** | Fwd | AGACTACAAAGGAGAGCAAACC |
| ***NM_008344.3*** | Rev | GAACAGGATTGGGCCGTATAG |
| ***Mmp9*** | Fwd | AAAGACCTGAAAACCTCCAACCT |
| ***NM_013599.4*** | Rev | TGTAACCATAGCGGTACAAGTATGC |
| ***Ngfr*** | Fwd | GTCCACACTCCTTCTCTTACAC |
| ***NM_033217.3*** | Rev | GTCCACACTCCTTCTCTTACAC |
| ***Rplp0 (m36B4)*** | Fwd | TCATCCAGCAGGTGTTTGACA |
| ***NM_007475.5*** | Rev | GGCACCGAGGCAACAGTT |
| ***Actb*** | Fwd | GGCTGTATTCCCCTCCATCG |
| ***XM_030254057.1*** | Rev | CCAGTTGGTAACAATGCCATGT |

### Table S4. siRNAs.

| Gene name | siRNA | Duplex Sequence (5'-3') |
|---|---|---|
| ***Mlana (mouse)*** ***Gene ID: 77836*** ***NM_029993.1*** | *siMlana-1* | rGrGrCrGrUrCrArUrArUrUrGrGrUrArUrUrCrArArArArAAC |
| | | rGrUrUrUrUrUrGrArArUrArCrCrArArUrArUrGrArCrGrCrCrUrU |
| | *siMlana-2* | rArCrGrArArGrUrGrGrArUrArCrArGrArArCrCrUrUrGrATG |
| | | rCrArUrCrArArGrGrUrUrCrUrGrUrArUrCrCrArCrUrUrCrGrUrCrU |
| ***Ngfr (mouse)*** ***Gene ID: 18053*** ***NM_033217.3*** | *siNgfr-1* | rCrCrArArCrArGrUrCrArGrArArCrCrGrArGrCrArUrCrUCT |
| | | rArGrArGrArUrGrCrUrCrGrGrUrUrCrUrGrArCrUrGrUrUrGrGrGrC |
| | *siNgfr-2* | rGrCrCrUrGrGrArCrArGrUrGrUrUrArCrGrUrUrCrUrCrUGA |
| | | rUrCrArGrArGrArArCrGrUrArArCrArCrUrGrUrCrCrArGrGrCrArG |
| ***Negative control*** | *siCtrl* | N/A (Proprietary, AllStar Neg. Control siRNA, QIAGEN #1027218) |

**Table S5. Detailed information of antibodies/dyes used for flow cytometry.**

| Antibodies/Dye | Company | Figure |
|---|---|---|
| Aqua Fixable Viability dye | Invitrogen | Figure 5A, S5A-C and S5E |
| BV785 – CD45 Clone: 30-F11 | BD Biosciences | |
| BV650 – CD3e Clone 145-2C11 | BD Biosciences | |
| PerCP-Cy5.5 – CD8a Clone: 53-6.7 | BD Biosciences | |
| PE-CF594 – CD31 Clone: MEC 13.3 | BD Biosciences | |
| e450 – CD11b Clone: M1/70 | eBioscience | |
| PE – F4/80 Clone: BM8 | eBioscience | |
| A488 – Gr-1 (Ly-6G) Clone: RB6-8C5 | eBioscience | |
| APC – Ly-6C Clone: AL-21 | BD Biosciences | |
| APC-e780 – CD11c Clone: N418 | eBioscience | |
| Aqua Fixable Viability dye | Invitrogen | Figure 5D, 5E, 5I, S5F-H |
| CFSE proliferation dye | Tonbo | |
| LEAF Purified anti-mouse CD3e Clone 145-2C11 | Biolegend | |
| PerCP-Cy5.5 – CD8a Clone: 53-6.7 | BD Bioscience | |
| APC-Cy7 CD4 Clone: RM4-5 | BD Bioscience | |
| PE IFNγ Clone: XMG1.2 | BD Bioscience | |
| Fixable Viability Dye eFluor 506 | eBioscience | Figure 7G |
| BUV737 – CD3 (clone 17A2) | BD Biosciences | |
| APC-Cy7 – CD45.2 (clone104) | eBioscience | |
| PE-Cy7 – F4/80 (clone BM8) | eBioscience | |
| PerCP-Cy5.5 – CD11c (clone HL3) | BD Biosciences | |
| APC – CD19 (clone eBio1D3) | eBioscience | |
| FITC – I-A[b] (clone AF6-120.1) | BD Biosciences | |
| Pacific Blue – CD11b (clone M1/70) | eBioscience | |
| PE- CD103 (clone 2E7) | eBioscience | |
| Fixable Viability Dye eFluor 780 | eBioscience | Figure S7F |
| BUV737 – CD3 (clone 17A2) | BD Biosciences | |
| Alexa Fluor 700 – CD4 (clone GK1.5) | ThermoFisher | |
| V500 – CD8 (clone 53-6.7) | BD Biosciences | |
| BV 650 – PD-1 (clone J43) | BD Biosciences | |
| PE-Cy7 – CD25 (clone PC61) | BD Biosciences | |
| PerCP-Cy5.5 – CD62L (clone MEL-14) | eBioscience | |
| APC – CD44 (clone IM7) | BD Biosciences | |
| FITC – Foxp3 (clone FJK16s) | eBioscience | |
| BUV395 – Ki67 (clone B56) | BD Biosciences | |
| Pacific Blue – Helios (clone 22F6) | BioLegend | |
