## Supplementary material for "The MNK1/2-eIF4E axis drives melanoma plasticity, progression, and resistance to immunotherapy": Supp. Fig. S1-S7

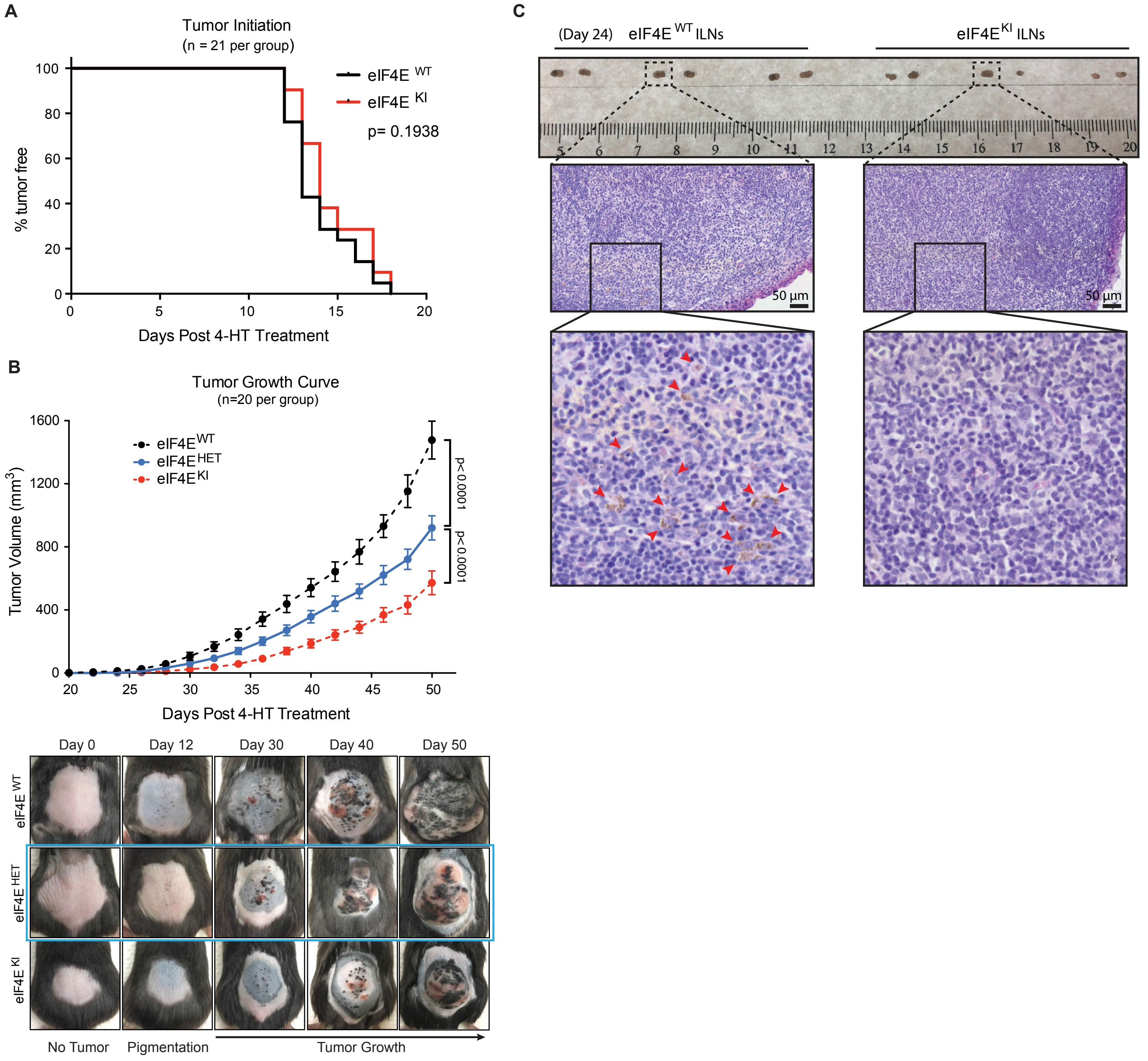


**Supplemental Figure S1. Impact of phospho-eIF4E-deficiency on melanoma outgrowth and metastasis.**
**(A)** Kaplan–Meier curves showing initiation of pigmented lesion on eIF4E$^{WT}$ and eIF4E$^{KI}$ mice (n = 21 per group). Log-rank test. **(B)** Tumor growth curve (top) and representative pictures (bottom) of eIF4E$^{HET}$ mice (n = 20) at indicated time points after 4-HT administration, comparing to eIF4E$^{WT}$ and eIF4E$^{KI}$ mice (as shown in Figure 1, n = 20 per group). Values are represented as mean ± SEM. Two-way ANOVA. **(C)** Pictures of ILNs (top) and representative images of H&E-stained ILN sections (bottom) of eIF4E$^{WT}$ and eIF4E$^{KI}$ (n = 3 per group) mice (Day 24). Early metastases are indicated with red arrows.

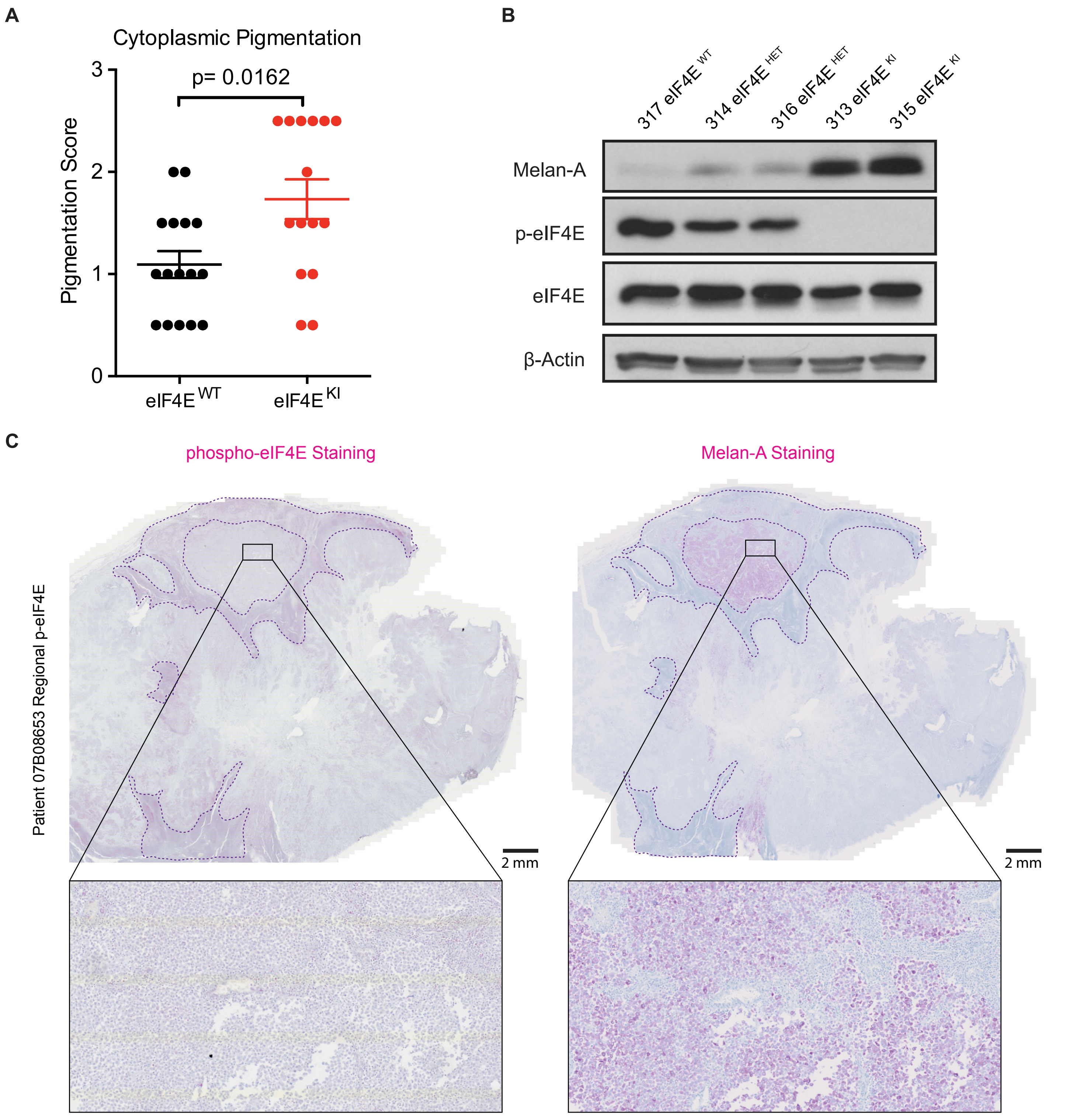


**Supplemental Figure S2. Phospho-eIF4E expression is negatively correlated with the expression of Melan-A in melanoma. (A)** Pigmentation score of $eIF4E^{WT}$ (n = 16) and $eIF4E^{KI}$ (n = 15) tumors (Day 50). Mann Whitney test. **(B)** Western blot of the indicated proteins in a cohort of $eIF4E^{WT}$, $eIF4E^{HET}$ and $eIF4E^{KI}$ primary melanoma samples. **(C)** IHC staining showing the expression of phospho-eIF4E and Melan-A in a patient derived melanoma sample with regional phospho-eIF4E expression.

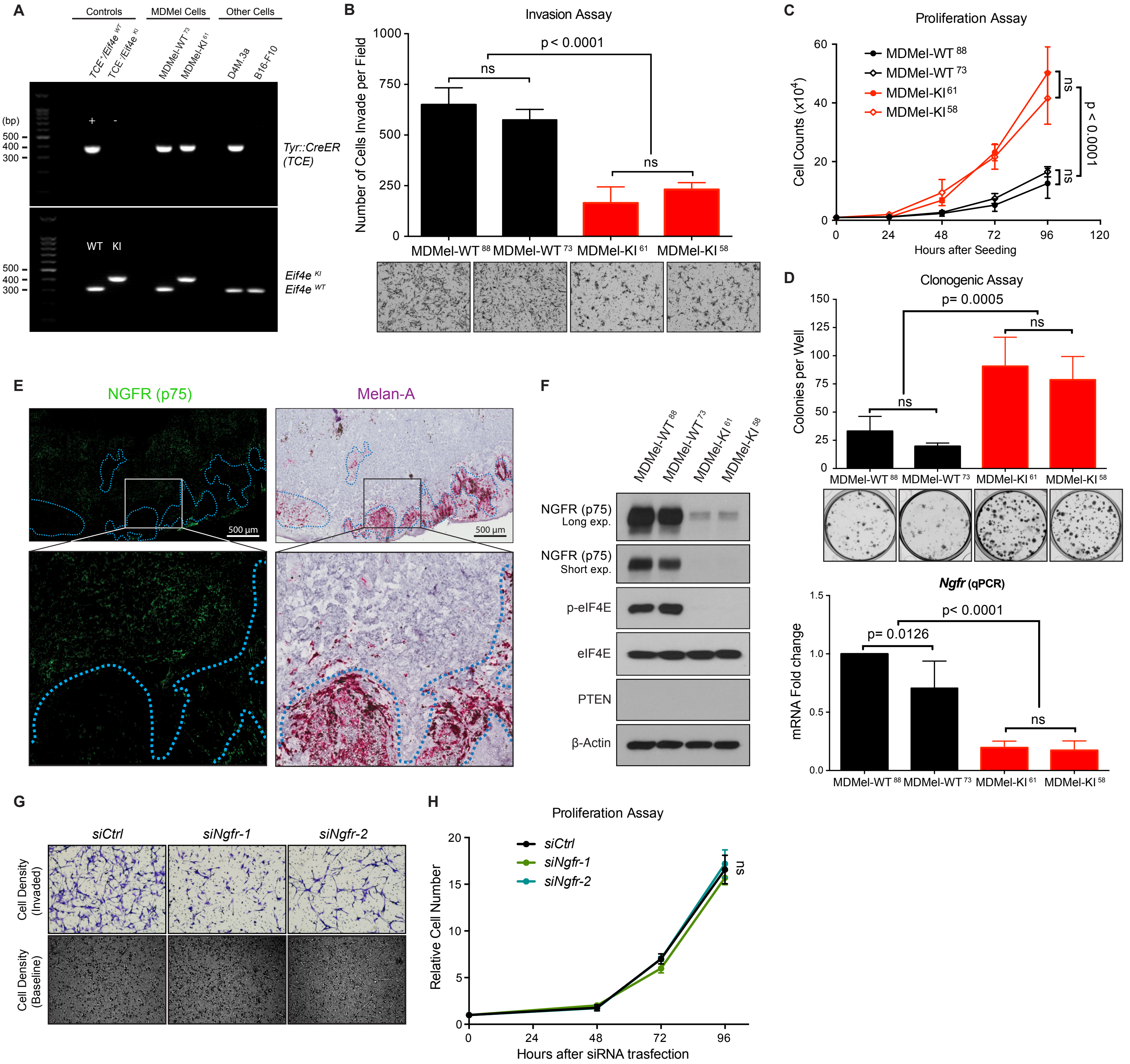

**Supplemental Figure S3. Phospho-eIF4E deficient melanoma cells exhibit less invasive phenotype with decreased NGFR expression.**
**(A)** Genotyping of murine melanoma cell lines MDMel-WT$^{73}$, MDMel-KI$^{61}$, D4M.3a and B16-F10 by Multiplex PCR. **(B-D)** Characterization of the invasion **(B)**, proliferation **(C)** and clonogenicity **(D)** of the murine melanoma cell lines MDMel-WT$^{88}$, MDMel -WT$^{73}$, MDMel-KI$^{61}$ and MDMel-KI$^{58}$ (n = 3 independent experiments). Two-way ANOVA with Sidak correction for multiple comparisons. **(E)** Representative images of an eIF4E$^{KI}$ melanoma sample (Day 50, from a total n = 8) stained for NGFR using IF (left) and for Melan-A using IHC (right). **(F)** Western blot analysis (left) and RT-qPCR analysis (right) assessing NGFR expression in the murine melanoma cell lines MDMel-WT$^{88}$, MDMel-WT$^{73}$, MDMel-KI$^{61}$ and MDMel-KI$^{58}$ (n = 3). Two-way ANOVA with Sidak correction for multiple comparisons. **(G)** Representative images with original magnification (×10) showing migrated (top panels) and baseline (bottom panels) MDMel-WT$^{73}$ cells that were silenced or not for *Ngfr*. **(H)** Proliferation of MDMel-WT$^{73}$ cells following *Ngfr* knockdown or control siRNA transfection. Two-way ANOVA with Sidak correction for multiple comparisons. All values are represented as mean ± SD.

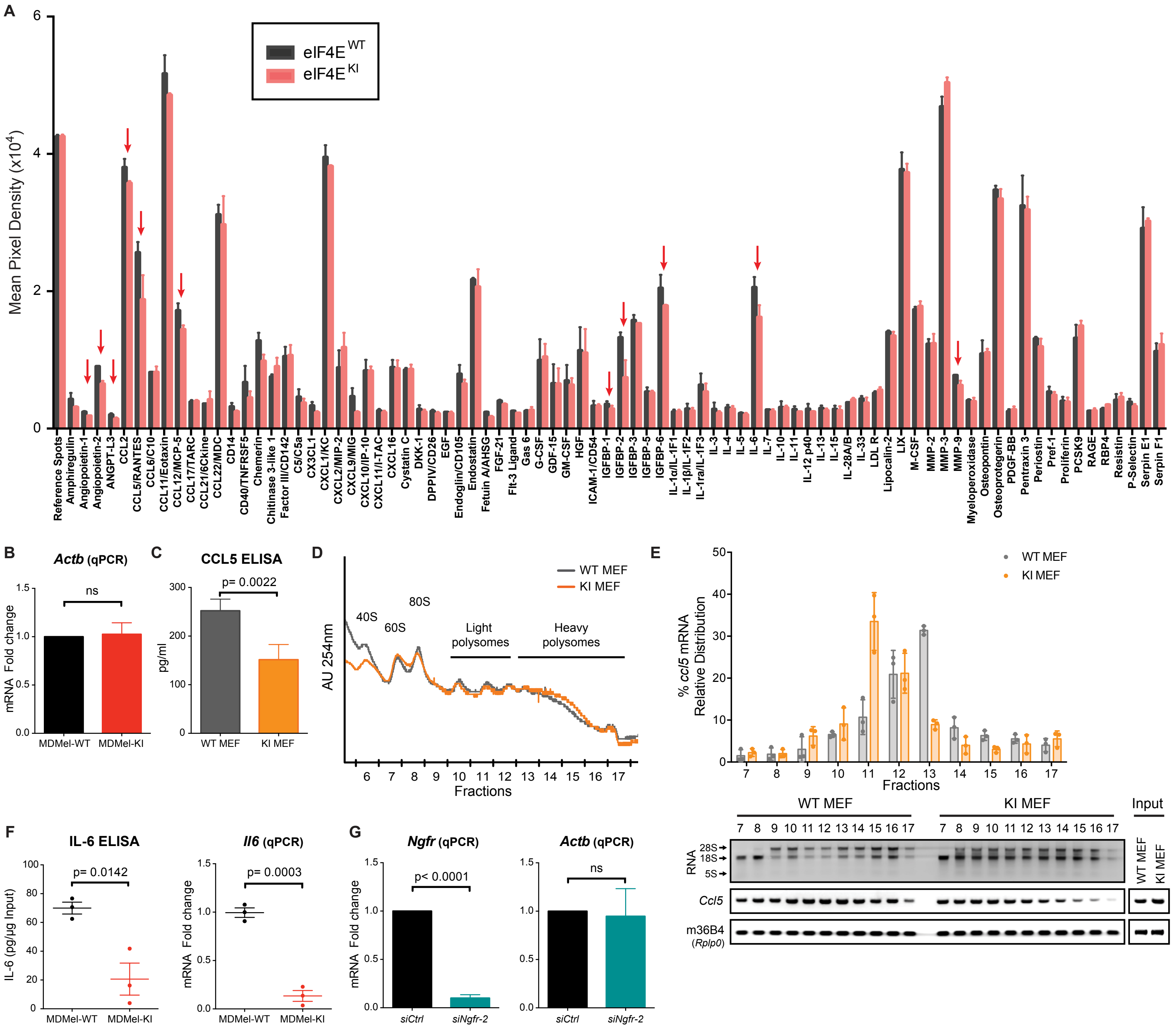


**Supplemental Figure S4. Phospho-eIF4E-defficiency impairs pro-inflammatory cytokine secretion by melanoma.**
**(A)** Cytokine/chemokine profiles from the conditioned medium of eIF4E$^{WT}$ and eIF4E$^{KI}$ primary melanoma cultures. Values represent the mean ± SEM (Day 35, n = 2 mice per genotype). **(B)** Percentage of housekeeping gene *Actb* mRNAs in MDMel-KI cells relative to MDMel-WT cells, normalized to m36B4 (*Rplp0*) as a reference gene (mean ± SD, n = 5). **(C)** CCL5 concentration in the conditioned medium of WT and KI MEFs. Values represent the mean ± SD (n=4). Two-sided unpaired t-test. **(D)** Polysome profiles of WT and KI MEFs (representative of n = 3). **(E)** Percentage of transcripts in each polysomal fraction from WT and KI MEFs, quantified by RT-qPCR (top) and representative image showing ribosomal RNAs and PCR-amplified cDNA fragments of indicated targets (bottom). Values represent the mean ± SD (n = 3). **(F)** IL-6 concentration in the conditioned medium (left) and relative *Il6* mRNA levels (right) from MDMel-WT and MDMel-KI derived melanomas (Day 22, n = 3 per group, see Figure 6A-C). Data are represented as mean ± SEM. Two-sided unpaired t-test. **(G)** Levels of *Ngfr* (left) and *Actb* (right) mRNA in *siNgfr-2*-transfected MDMel-WT cells relative to the control group, normalized to m36B4 (*Rplp0*) as a reference gene. Values represent the mean ± SD (n = 3). Two-sided unpaired t-test.

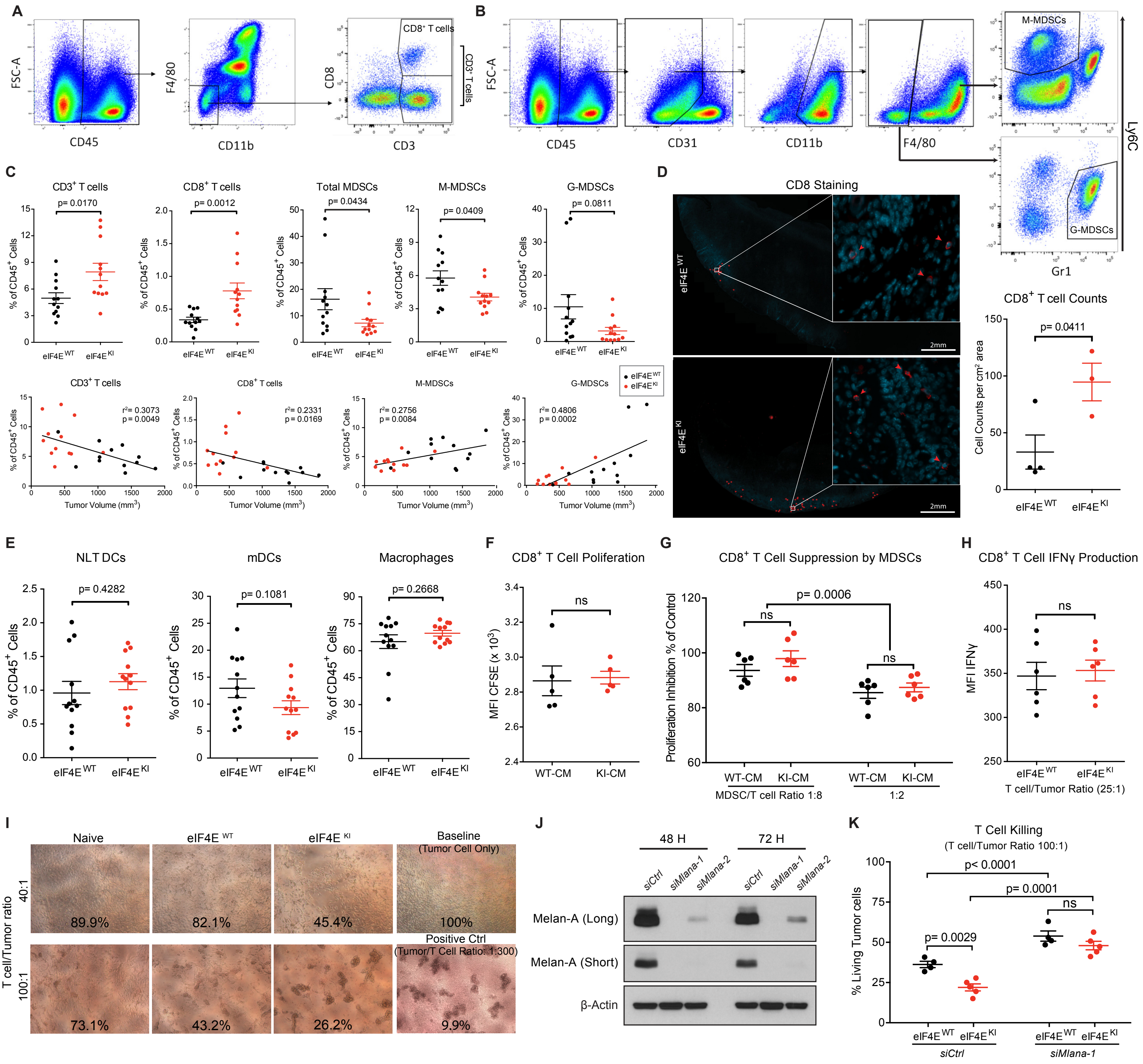


**Supplemental Figure S5. Characterizing the impact of phospho-eIF4E in melanoma immunosuppression.**
**(A and B)** Representative gating strategy to identify populations of T cells **(A)** and MDSCs **(B)**. Cells were pre-gated for viable, single cells. **(C)** Percentages of indicated tumor-infiltrating immune cells out of total CD45$^+$ cells (top, two-sided unpaired t-test) and their correlations with the size of the corresponding tumors (bottom, Pearson correlation coefficient) (n = 12 per group). **(D)** Representative images show the expression of CD8 via IF staining in eIF4E$^{WT}$ (n = 4) and eIF4E$^{KI}$ (n = 3) melanomas (left), and graphed are the CD8$^+$ cell counts relative to tumor size (right). Two-sided unpaired t-test. **(E)** Percentages of indicated tumor-infiltrating immune cells out of total CD45$^+$ cells in eIF4E$^{WT}$ and eIF4E$^{KI}$ tumors (n = 12 per group). Two-sided unpaired t-test. **(F)** Mean fluorescence intensity (MFI) of CFSE in CD8$^+$ T cells following a 72-hour CD3/CD28 stimulation in the conditioned medium from eIF4E$^{WT}$ and eIF4E$^{KI}$ primary melanoma cultures (WT-CM/KI-CM, n = 5 mice per group). Two-sided paired t-test. **(G)** Inhibition of CD8$^+$ T-cell proliferation by MDSCs at the indicated ratio in WT-CM or KI-CM (n = 6 per group), relative to the corresponding MDSC-free control group. Two-way RM ANOVA with Sidak correction for multiple comparisons. **(H)** MFI of IFNγ in CD8$^+$ T cells isolated from ILNs of eIF4E$^{WT}$ and eIF4E$^{KI}$ tumor-bearing mice (n = 6 per group), co-cultured with B16-F10 cells at indicated ratio. Two-sided paired t-test. **(I)** Representative images of B16-F10 cells co-cultured with T cells from eIF4E$^{WT}$, eIF4E$^{KI}$ tumor-bearing animals (n = 4 per group) and non-tumor-bearing mice (n = 2) at indicated ratio. Percent viability of tumor cells in each condition, relative to the baseline control, is indicated. **(J)** Western blot of Melan-A in B16-F10 cells after transfection with *siMlana-1*, *siMlana-2* (see Table S4) and control siRNA at indicated time points. **(K)** Percent viability of B16-F10 cells co-cultured with T cells from eIF4E$^{WT}$ (n = 4) and eIF4E$^{KI}$ tumor-bearing animals (n = 5) upon Melan-A knockdown (*siMlana-1*), relative to corresponding control groups. Two-way RM ANOVA with Sidak correction for multiple comparisons. For *in vivo* experiments **(A-E)**, all tumors were resected on Day 50. For *ex vivo* assays **(F-K)**, all tumor-bearing mice were sacrificed between Day 35 and Day 38. All values are represented as mean ± SEM.

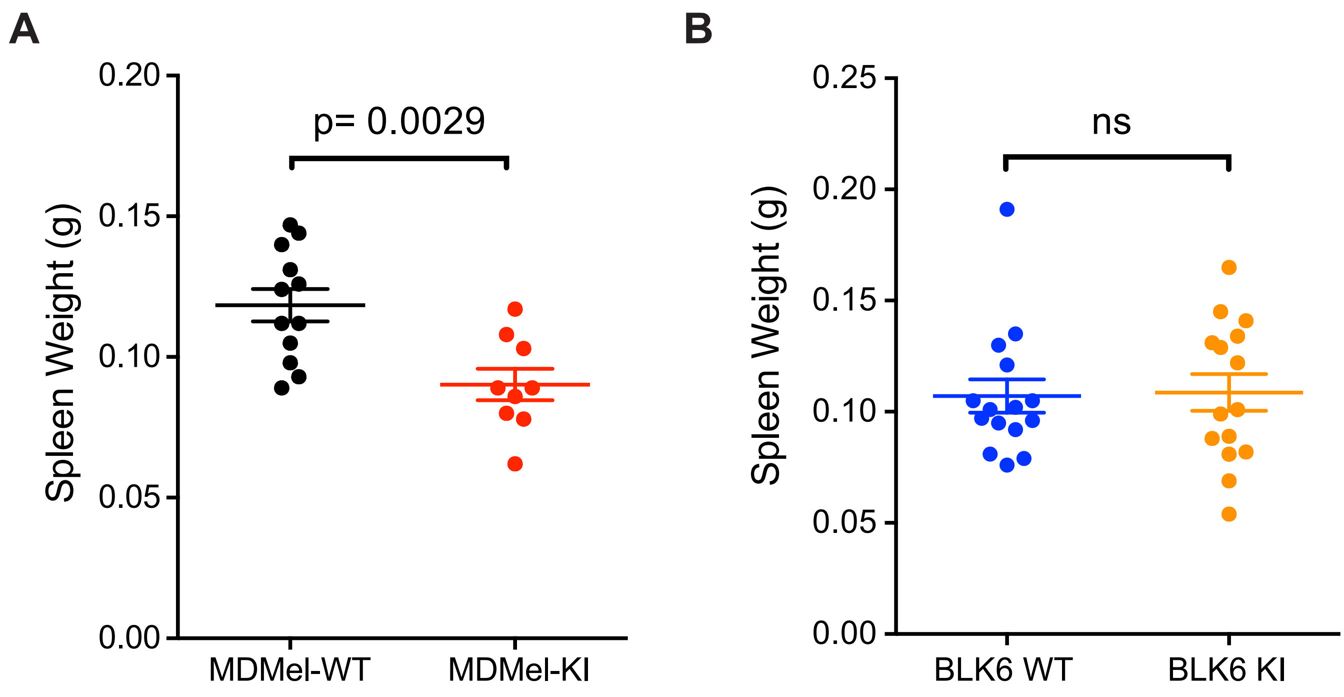


**Supplemental Figure S6. Dissecting tumor cell-intrinsic and -extrinsic roles of phospho-eIF4E using syngeneic melanoma models.**
**(A and B)** Spleen weight (Day 22) of BLK6 WT mice bearing MDMel-WT (n = 12) and MDMel-KI (n = 9) derived melanomas **(A)**, and spleen weight (Day 21) of BLK6 WT and BLK6 KI mice (n = 15 per genotype) bearing D4M.3a-derived melanomas **(B)**. Two-sided unpaired t test. All values are represented as mean ± SEM.

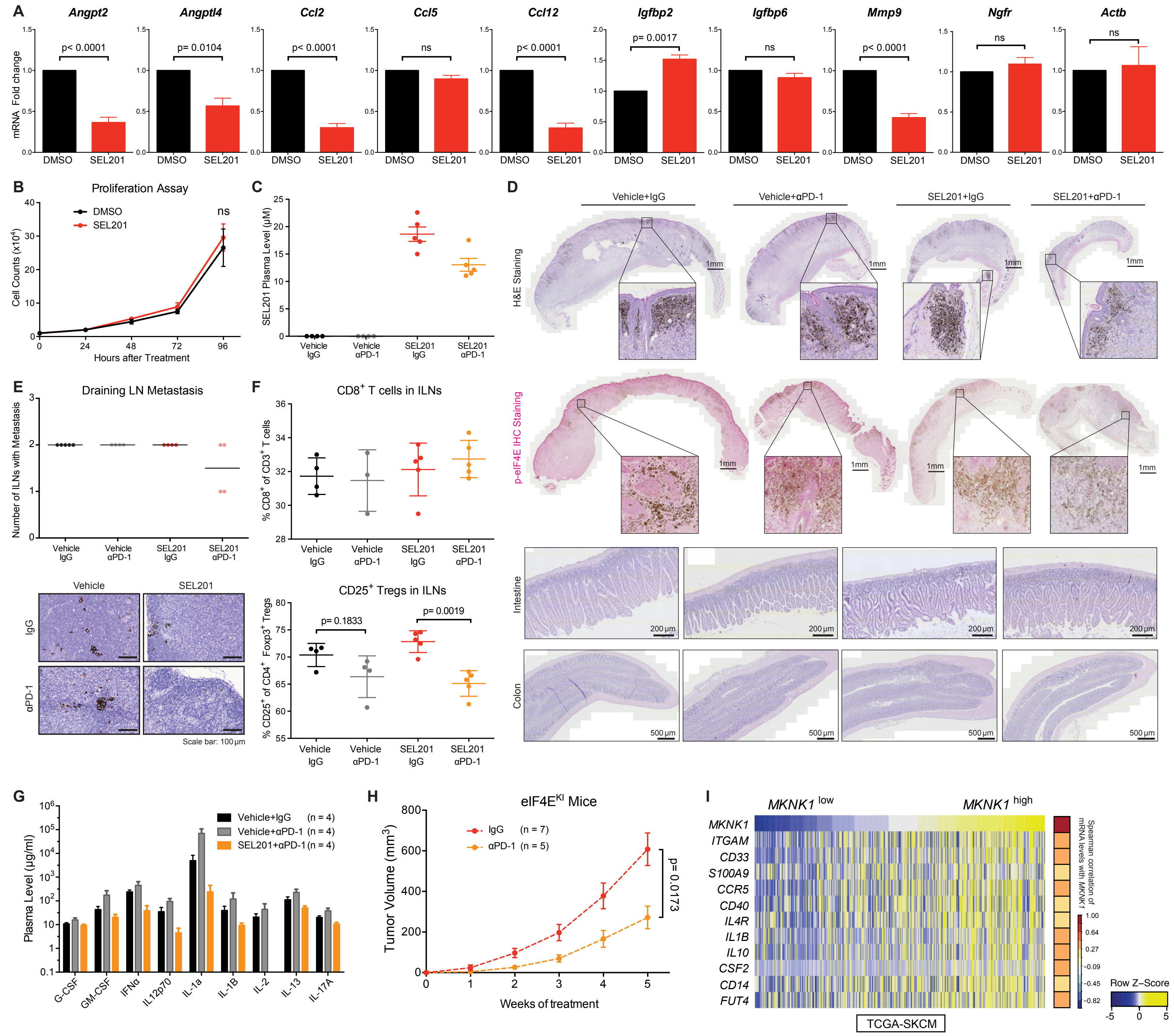


**Supplemental Figure S7. Evaluating the efficacy of blocking phospho-eIF4E using the MNK1/2 inhibitor SEL201, in combination with the anti-PD-1 immunotherapy in melanoma.**

**(A and B)** Characterization of melanoma cells treated with SEL201 (2.5 µM) *in vitro*. **(A)** Percentage of the indicated mRNAs in SEL201-treated MDMel-WT cells relative to the DMSO-treated control group, normalized to m36B4 (Rplp0) as a reference gene (n = 3 for *Angptl4*, *Ccl5*, *Igfbp2* and *Igfbp6*, n = 4 for the rest). Two-sided unpaired t-test. **(B)** Proliferation of MDMel-WT cells with treated with SEL201 or DMSO as control. Two-way ANOVA with Sidak correction for multiple comparisons. All *in vitro* data are represented as mean ± SD. **(C-H)** Evaluation of SEL201 treatment in combination with anti-PD-1 monoclonal antibody (αPD-1) *in vivo*. **(C)** SEL201 level detected in the plasma harvested from mice treated with the indicated compound. **(D)** Representative images of melanomas: H&E-staining (top), IHC staining for phospho-eIF4E (middle), and H&E-stained intestine and colon samples from each group (n = 14 per group). **(E)** Numbers of metastasis-positive ILNs per mouse (top) and representative images of H&E-stained ILNs (bottom) are presented for each group. **(F)** Percentage of CD8$^+$ T cells out of total CD3$^+$ T cells (top) and percentage of CD25$^+$ Tregs of the total CD4$^+$ Foxp3$^+$ Tregs (bottom) in ILNs from animals in each group. Two-way ANOVA, Tukey's multiple comparisons test. **(G)** Concentrations of toxicity-related growth factors/cytokines/chemokines in control (Vehicle+IgG), anti-PD-1 alone (Vehicle+αPD-1) and combination therapy (SEL201+αPD-1) groups. Plasma concentrations of secreted factors were detected by the MAGPIX multiplexing system. **(H)** Tumor growth curve of eIF4E$^{KI}$ melanoma-bearing mice administered IgG or αPD-1. Two-way ANOVA with Sidak correction for multiple comparisons. **(I)** Expression of *MKNK1* and genes indicating MDSC infiltration (HTSeq - FPKM) in GDC TCGA Melanoma dataset (SKCM, n = 472). Spearman rank-order. For *in vivo* experiments **(C-H)**, all values represent the mean ± SEM, with numbers of animals in each group (minimum n = 4) indicated in the figure.
